# Modeling the Memory of Unmyelinated Axons: Integration of a Data-Driven Approach with Physiological Memory Concept

**DOI:** 10.1101/2025.09.24.678232

**Authors:** Anna Maxion, Jenny Tigerholm, Barbara Namer, Ekaterina Kutafina

**Affiliations:** Joint Research Center for Computational Biomedicine, Medical Faculty, RWTH Aachen University, 52072 Aachen, Germany; Scientific Center for Neuropathic Pain Aachen, SCN Aachen, 52072 Aachen, Germany; Department of Anesthesiology, Intensive Care, Emergency and Pain Medicine, University Hospital Würzburg, Center for Interdisciplinary Pain Medicine, 97080 Würzburg, Germany; Institute for Biomedical Informatics, Faculty of Medicine, University Hospital Cologne, University of Cologne, 50937 Cologne, Germany

## Abstract

In this study, we present a simplified and resource-efficient computational model to predict activity-dependent conduction velocity changes of action in unmyelinated axons. This model serves as a complementary tool to Hodgkin-Huxley models. Our approach is based on the concept of "memory," where the speed of subsequent action potentials is modulated by prior activity. We utilized microneurography data from 95 mechano-insensitive C-fibers of healthy human participants, including both sexes, across various stimulation protocols to optimize model parameters. The model incorporates linear long-term and non-linear short-term memory components. By convolving the history of recorded action potentials with the memory function, the model can effectively predict the propagation speed of subsequent action potentials with low mean squared errors for the proposed one-dimensional and two-dimensional memory functions. This computational framework provides insights into the dynamics of unmyelinated axons under varying conditions and thus in signal processing along the axon and the short-term memory of axons. The model’s rapid computation times make it suitable for real-time applications in electrophysiological experiments.

**Significance Statement:** This study introduces a novel model for simulating activity-dependent conduction velocity changes in unmyelinated axons, which crucially shape signal processing during conduction. While traditional Hodgkin-Huxley (HH) models offer valuable physiological insights, they are computationally intensive and challenging to adapt due to numerous parameters. Our approach leverages the concept of fiber “memory,” capturing how prior activity shapes conduction. It requires fewer parameters and therefore makes it easier to fit to diverse datasets, including patient data. Importantly, our model is highly efficient and suitable for real-time applications, enabling rapid simulation and analysis that are not feasible with HH models.

## 1. Introduction

Understanding the function and dynamics of nerve fibers is fundamental to advancing our knowledge of the nervous system and developing treatments for neurological conditions. Unmyelinated axons play a crucial role in transmitting sensory signals and particularly in nociception and pain signaling (Dubin and Patapoutian, 2010). They are also abundantly present in the central nervous system (Soleng et al., 2003; Westenbroek et al., 1989).

An important aspect of the physiology of unmyelinated axons is characterized by activity-dependent conduction velocity changes, such as activity-dependent slowing (ADS) during repeated stimulation (Carr, 2013; Schmelz et al., 1995). ADS can be used to differentiate between different C-fiber subtypes that exhibit characteristic conduction velocity changes in response to stimulation (Obreja et al., 2010; Serra et al., 1999; Weidner et al., 1999). The magnitude of ADS depends on several factors, including frequency, duration, and intensity of previous activation of the axon (Ackerley and Watkins, 2018; Namer and Lampert, 2025; Weidner et al., 2000). When a fiber has already undergone substantial conduction velocity slowing, e.g. after prolonged activation with a high action potential count, additional stimulation can lead to a relative increase in conduction velocity rather than further slowing (Weidner et al., 2002), revealing complex adaptation dynamics depending on prior activity. Those adaptations lead to either increased frequencies or decreased frequencies arriving at the spinal cord and thereby provide a filter function of the axon cutting out medium frequencies leading to a more bursting behaviour (Bucher and Goaillard, 2011).

Abnormal ADS patterns have been observed in inflammatory pain, neuropathic pain and small fiber neuropathies (Dickie et al., 2017; Kist et al., 2016; Kleggetveit et al., 2012; Namer et al., 2015a; Ørstavik et al., 2006, 2003), making it a valuable marker for detecting and characterizing nerve dysfunction.

To study activity-dependent conduction velocity changes, microneurography is a valuable technique that allows direct recording of nerve fiber activity in awake humans (Ackerley and Watkins, 2018; Namer and Lampert, 2025; Schmelz and Schmidt, 2010; Vallbo, 2018). However, microneurography is a tedious and time-consuming procedure, requiring long experimental sessions to obtain sufficient data for analysis.

Computational models offer an alternative approach by simulating nerve fiber behavior under different conditions. Complex multicompartmental models, including Hodgkin-Huxley (HH) models, incorporate detailed biophysical dynamics of ionic currents and membrane potentials (Beeman, 2022; He et al., 2020; Petersson et al., 2014; Tigerholm et al., 2014). In our work (Maxion et al., 2023), we showed the efficiency of the HH approach to assess altered molecular mechanisms in patients. However, this level of detail often results in long computation times, high computational resource demands and a complex parameter fitting process.

In our other work (Kutafina et al., 2022), we investigated a fully data-driven approach to modelling of spike timing, through relating the spiking history of the fiber to the magnitudes of activity-dependent conduction velocity changes. This approach showed computational efficiency, however, the physiological knowledge provided by the HH approach was missing. Here, we present a new model that balances the physiological sophistication of the HH model and the computational efficiency of a purely data-driven approach by explicitly incorporating “memory” of unmyelinated axons: the influence of past activity on current conduction latency using a convolution-based approach inspired by geophysical wave propagation models (N. van Dalen et al., 2013).

The proposed model is designed for easy adaptation and predictive modeling, enabling simulations of high-frequency stimulation protocols. It could aid in spike sorting and predict spike times in real-time or during post-experiment analysis, facilitating longer axon simulations. The study aims to create a simple computational model that efficiently predicts action potential propagation latency and provides insights into the complex memory function of unmyelinated axons. This model has significant implications for pain research, as it can enhance our understanding of nociceptive signaling mechanisms. Additionally, it may contribute to a broader comprehension of central nervous system functions, shedding light on how neural dynamics influence various physiological processes.

## 2. Materials and Methods

We present a computational model developed to simulate axonal memory and predict changes in action potential latencies in response to the spiking history. The model operates by taking an input spike train and applying a predefined memory function to adjust the latencies of subsequent spikes as they travel along the axon (Figure 1). This transformation reflects the influence of prior activity on the timing of action potentials. The input spike train could also be interpreted as a stimulation signal, assuming each stimulation pulse elicits exactly one action potential and spontaneous activity is absent, which is characteristic behavior observed in healthy C-nociceptors.

**Figure 1:**
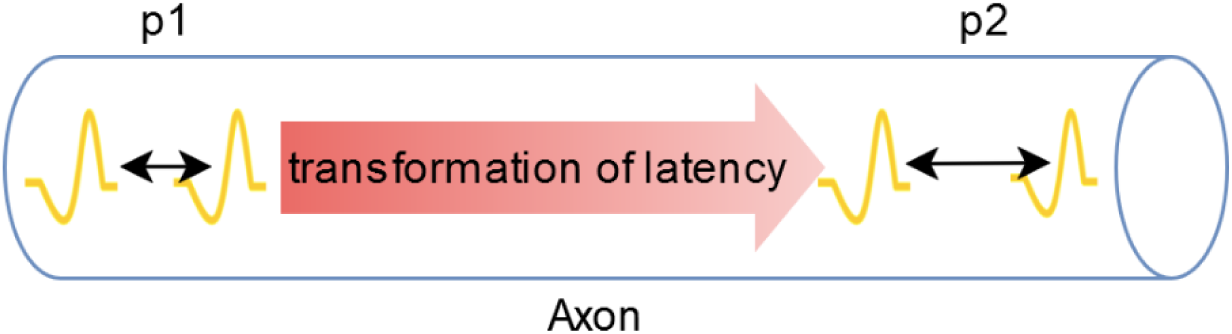
Underlying concept of the model: The spike train generated at point p1 of the axon (e.g., in response to electrical stimuli) propagates along the axon to the measurement point p2. During this transmission, the membrane properties are dynamically altered by preceding spikes. Consequently, the travel times between p1 and p2 vary for each spike, leading to modulation of the temporal pattern of the original spike train.

The subsequent subsections detail the concept and modelling of axonal memory through recovery cycles, the modeling of spiking history and the implementation of convolution techniques to capture the dynamic responses of unmyelinated axons. Further, it is described what data was used to optimize and validate the model, as well as the optimization technique.

### 2.1. Design of the Computational Model

The computational model employs a mathematical function that represents axonal memory, which is convolved with a defined discrete signal that represents the history of previous spikes. This convolution allows for the prediction of relative latencies based on the activity history of the fibers. The model was implemented in Python, leveraging its computational capabilities to efficiently calculate the convolution and test various memory functions. This implementation provides a flexible framework for refining the model and adapting it to experimental data.

#### 2.1.1 Concept of Axonal Memory using Recovery Cycles

Unmyelinated axons exhibit activity-dependent modulation of conduction velocity, which can be interpreted as a form of “memory”. This means that the conduction velocity at a given moment is influenced by the fiber’s previous activity. This behavior can be modeled using a memory function that reflects the fiber’s recovery dynamics.

A key example of this concept is provided by (Weidner et al., 2002), which illustrates the presumed time course of post-excitatory effects in human C-fibers (Figure 2). The figure summarizes how pre-existing conduction velocity slowing, expressed as the percentage decrease in conduction velocity, and the interstimulus interval following the last stimulation pulse influence the conduction velocity of the subsequent action potential. The key observations include the occurrence of supernormality, characterized by the increased conduction velocity, at high pre-existing slowing and short interstimulus intervals, while subnormality, or decreased conduction velocity, is observed at low pre-existing slowing. For longer interstimulus intervals, the effect on conduction velocity diminishes progressively, becoming negligible over time. Figure 8 depicts a similarly shaped model of memory function, constructed for the purpose of our study.

**Figure 2:**
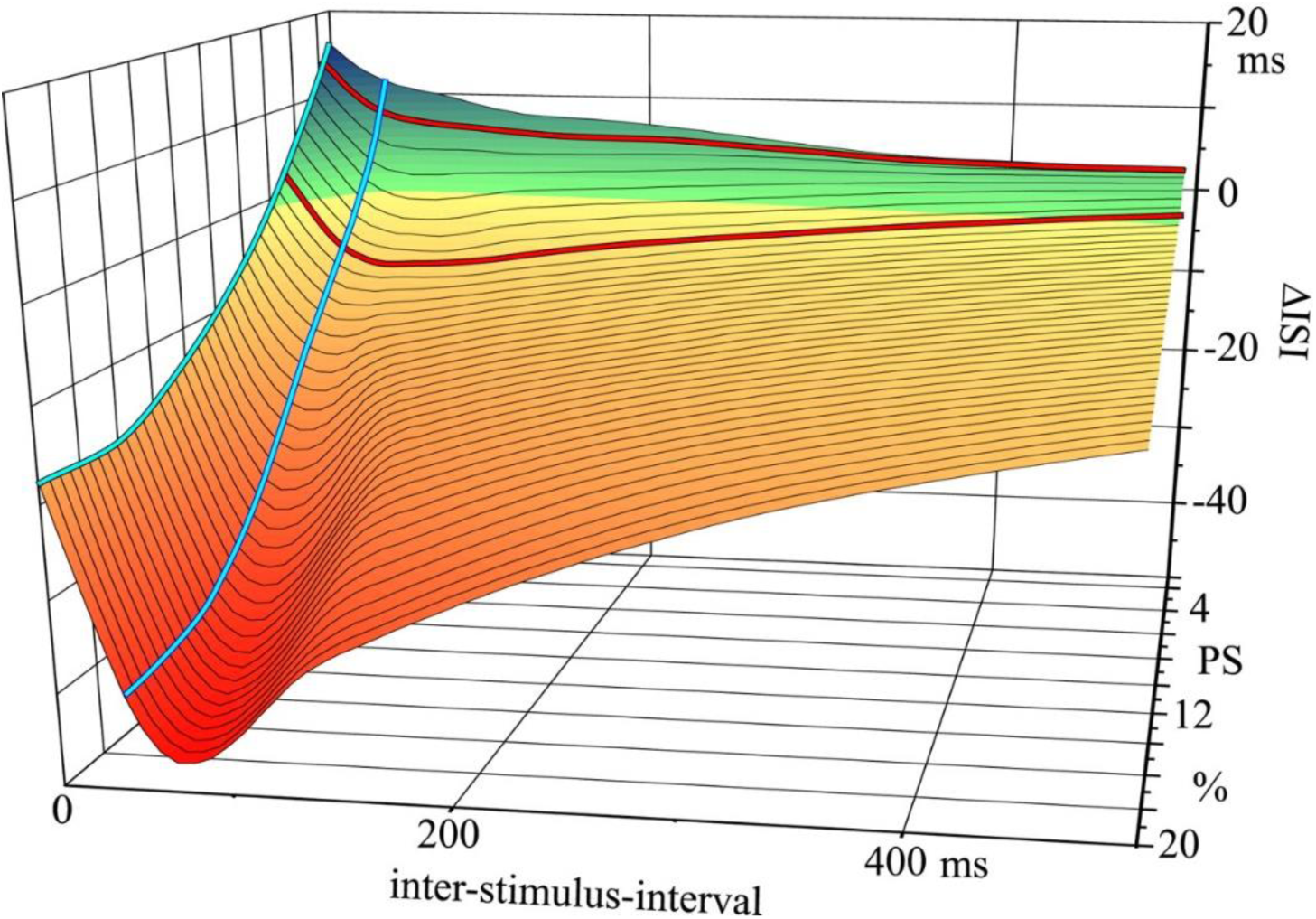
Presumed time course of post-excitatory effects in human C-fibers. This three-dimensional figure summarizes the effects of preexisiting slowing (*PS*, % decrease of conduction velocity) and interstimulus interval on ΔISI (supernormality in *yellow-red* and subnormality in *green*). Supernormality of conduction is most pronounced at intense PS and short interstimulus intervals, whereas subnormality is found at a low degree of PS. *Blue lines* represent findings from (Weidner et al., 2002). Data for the *red lines* is taken from (Weidner et al., 2000a). Figure and legend text from (Weidner et al., 2002), Copyright 2002 Society for Neuroscience.

Pre-existing slowing serves as a measure of previous activity (PA) within the fiber: generally, increased PA results in more conduction velocity slowing. Additionally, the interstimulus interval could also be interpreted as an interspike interval (ISI), when we assume that each stimulus elicits exactly one spike and there is no spontaneous activity.

To translate these findings into a functional model, we defined a memory function that depends on ISI and/or PA. For long ISIs, the effect decreases linearly, while for short ISIs, supernormality occurs. This means the model axon has a non-linear short-term memory and a linear long-term memory. In the two-dimensional model, the transition between sub- and supernormality is influenced by the PA value. This results in a piecewise function m(t) that captures both the activity-dependent modulation of conduction velocity and its temporal recovery dynamics. The model is described in detail in the next sections.

#### 2.1.2. Modelling Spiking History

To model the previous spiking activity, we define a discrete signal function *s(t)* that assigns a value of 1 at specific points in time corresponding to the occurrence of spikes and 0 elsewhere. Mathematically, this is expressed as:

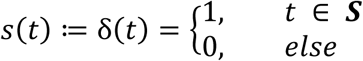

where ***S*** represents the set of times at which spikes occur.

#### 2.1.3. Convolution Model

To model the influence of previous spiking activity on the fiber’s latency, the signal *s(t)* is convolved with the memory function *m(t)*. The convolution operation is defined as:

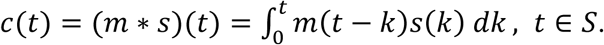

In this equation, *m(t)* represents the fiber’s memory dynamics, which encodes how previous activity affects the conduction latency at time *t* (see 2.2 for details). The convolution operation calculates the cumulative effect of all preceding activity on the latency.

At each time point *t*, the function *m(t−k)* determines the weight of past events, meaning how strongly a spike occurring at time *k* contributes to the fiber’s state at *t*. The signal *s(k)* indicates whether a spike occurred at time *k*, and its interaction with *m(t−k)* defines the effect of that spike on the fiber’s conduction velocity. Integrating over all past time points combines the effects of all previous spikes, yielding the total memory-driven modulation of conduction latency at *t*.

Importantly, the conduction latency *c(t)* is only calculated for *t* ∈ *S*, i.e., at time points when spikes occur. This ensures that the latency is updated only in response to actual spiking events, rather than continuously evolving in time.

### 2.2. Specific Memory Functions

The memory function is defined as a piecewise function that exhibits different behaviors depending on the length of the interspike intervals (ISIs). For ISIs greater than or equal to 0.5 seconds, we employed a linear memory function reflecting a linear decrease in conduction velocity for larger ISIs. In contrast, for ISIs shorter than 0.5 seconds, the function reflects supernormality, a phenomenon where conduction velocity temporarily increases following stimulation. The threshold of 0.5 seconds marks the point at which the supernormal phase in C-fibers concludes across various conditions and background frequencies. Beyond this interval, the fiber’s response becomes linear (see (Bostock et al., 2003)). We implemented two specific memory functions: a one-dimensional function that only depends on the ISI and a two-dimensional function that depends on the ISI and the pre-existing activity state (PA).

#### 2.2.1 One-Dimensional Memory Function

The proposed memory function is only dependent on the ISI, meaning that the effect of previous activity on conduction latency is determined by the time elapsed since the last spike.

The function is defined as:

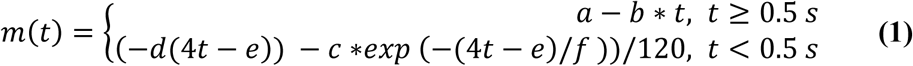

This formulation describes how the fiber "remembers" past activity and how its conduction latency changes over time.

For longer ISIs (*t≥0.5 s*), the memory function follows a linear decrease with increasing ISI. This describes that the influence of previous activity gradually fades over time, leading to a progressive normalization of conduction velocity.

For shorter ISIs (*t<0.5 s*), the function is exponential, capturing the rapid and transient effects of recent stimulation. The term *exp* (−*d*(4*t* − *e*)) represents an overall exponential decline, and the term *c* ∗*exp* (−(4*t* − *e*)/*f*)) defines the supernormal phase. The parameter *e* moves the function on the x-axis, d determines the steepness of the beginning of the function, f determines the width of the supernormal phase and *c* the depth of the supernormal phase (Figure 3).

**Figure 3:**
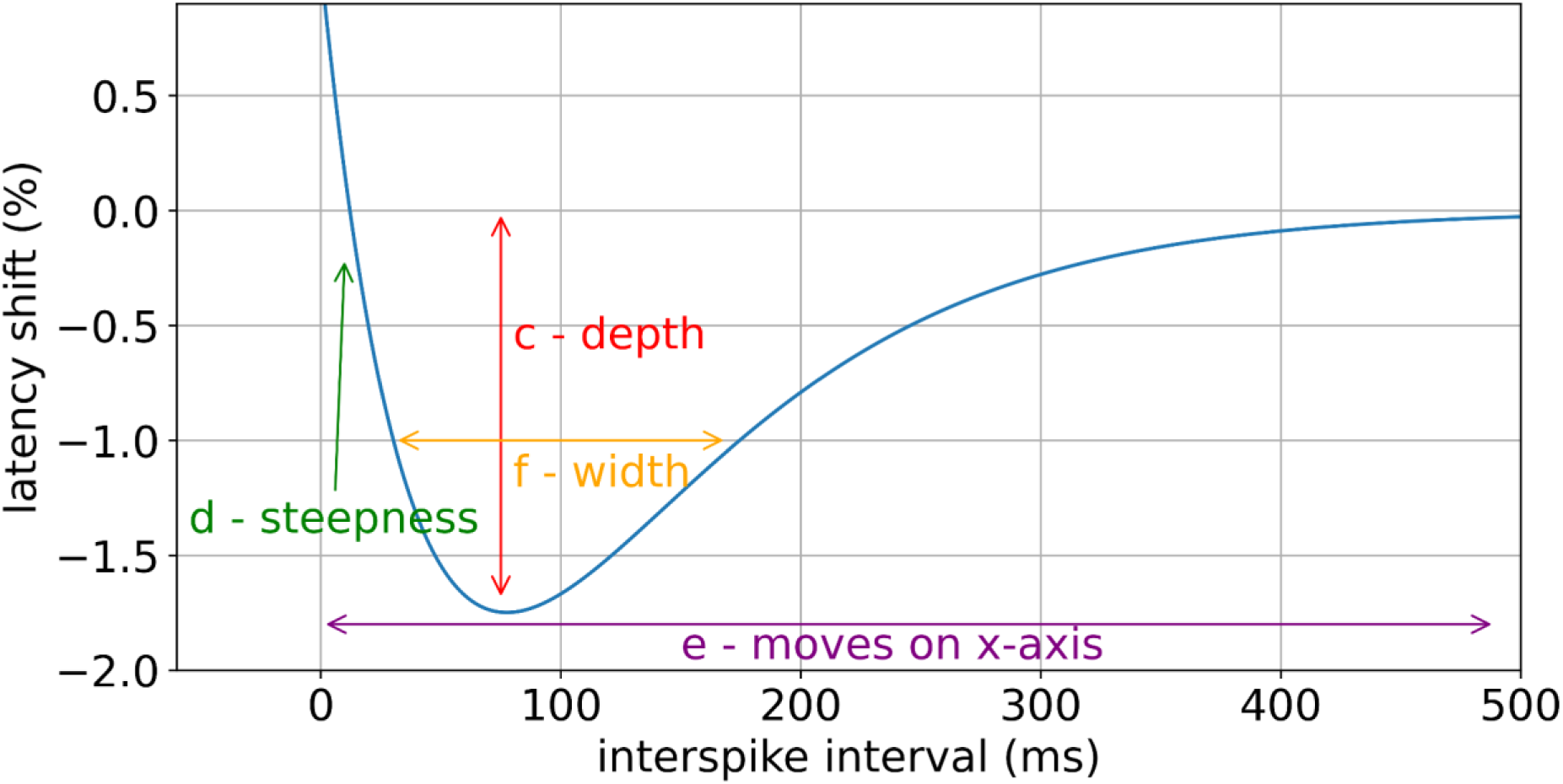
Visualization of parameters of non-linear memory function: The parameters explaining this function are highlighted with arrows in different colors. The parameter c controls the depth of the supernormal phase. *d* controls the steepness of the beginning of the curve. The constant *e* moves the function along the x-axis, determining the position of the supernormal phase. Parameter *f* controls the width of the supernormal phase.

#### 2.2.2 Two-Dimensional Memory Function

Since the memory function is not only dependent on ISI (*x*), but also on the pre-existing activity state (PA, *y*), we add this complexity level to the model and propose the two-dimensional modification of the memory function. It is defined as:

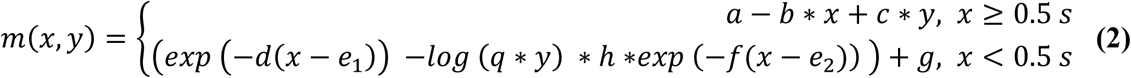

In the linear part of the function for larger ISIs (*x* ≥ 0.5 *s*), an additional term reflecting the dependence on PA has been incorporated as (*c* ∗ *y*).

For smaller ISIs (*x* < 0.5 *s*), the following modifications were made: the parameter *e* was divided into two distinct parameters *e*_1_ and *e*_2_, allowing independent movement of different parts of the function along the x-axis. The parameter c was replaced with the term *log* (*q* ∗ *y*) ∗ ℎ, which makes the depth of the supernormal phase dependent on the PA. Thus, larger values of PA result in a more pronounced supernormal phase. Additionally, the parameter g was introduced to be able to move the function on the vertical axis.

During the calculation of the convolution, the pre-existing activity state (*y*) is dependent on *x* and the set of stimulation pulses *S* = {*s*_1_, *s*_2_,…, *s*_*n*_}, *n* ∈ *N* and is defined as follows:

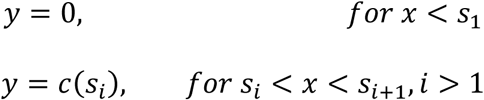

Before the first stimulation pulse is applied, there is no pre-existing activity. Hence, the fiber remains in its original state. Subsequently, the pre-existing activity state is defined as prior latency. Therefore, the memory function is technically only dependent on ISI (*x*) and the same convolution operation can be conducted as before.

### 2.3. Overview of Experimental Data

#### 2.3.1 Microneurography Methodology

Microneurography is a technique to record nerve signals from individual C-fibers. To achieve this, an uninsulated microelectrode with a tip diameter of 0.005 mm (Microneurography needle, FHC, Bowdoinham, US) is inserted into a fascicle of the superficial peroneal nerve at the ankle level, while a reference electrode is positioned superficially in the nearby skin.

Electrical pulses given by an insulated constant current stimulator (Digitimer DS7, Digitimer, Hertfordshire, UK), are applied to the receptive field of the fibers using superficially inserted electrodes (microneurography reference electrodes). This stimulation elicits action potentials in the terminal tree of the nerve fibers in the skin at a consistent latency, enabling the identification of multiple fibers within the same recording session. The mechanical thresholds of individual fibers are assessed using calibrated von Frey hairs and the marking method (Schmidt et al., 1995; Torebjörk and Hallin, 1974). Both the stimulation and the recording of fiber activity are performed using DAPSYS (Data Acquisition Processor System, Brian Turnquist, http://dapsys.net), which allows for real-time data acquisition and processing.

#### 2.3.2 Experimental Microneurography Data Obtained in Namer’s Lab

Microneurography data from 97 mechano-insensitive C-fibers of healthy individuals were used to optimize the computational model. Two fibers were excluded from this dataset: one due to incomplete data caused by recording issues and another because of misclassification in the original dataset. This resulted in a final dataset of 95 fibers.

The study adhered to the principles outlined in the Declaration of Helsinki and the statutory requirements of Germany. Ethical approval was obtained from the local ethics committees in Aachen (EK 141/19) and Erlangen (4361). All participants provided written informed consent to take part in the study.

For experimental stimulation, each fiber was subjected to electrical rectangular pulses with a duration of 0.5 ms. Prior to each protocol, electrical stimulation was paused for 2 minutes to ensure baseline stability.

Two distinct stimulation protocols were employed:

1. The first protocol involved a series of rising pulse frequencies designed for fiber type discrimination, termed the electrical identification (ELID) protocol. This consisted of 20 pulses at 1/8 Hz, followed by 20 pulses at ¼ Hz, then 30 pulses at ½ Hz, and concluding with 10 pulses at ¼ Hz (Figure 4A).
2. In the second protocol, each fiber was stimulated continuously at a frequency of 2 Hz for a duration of three minutes before returning to a frequency of ¼ Hz (Figure 4B).

**Figure 4:**
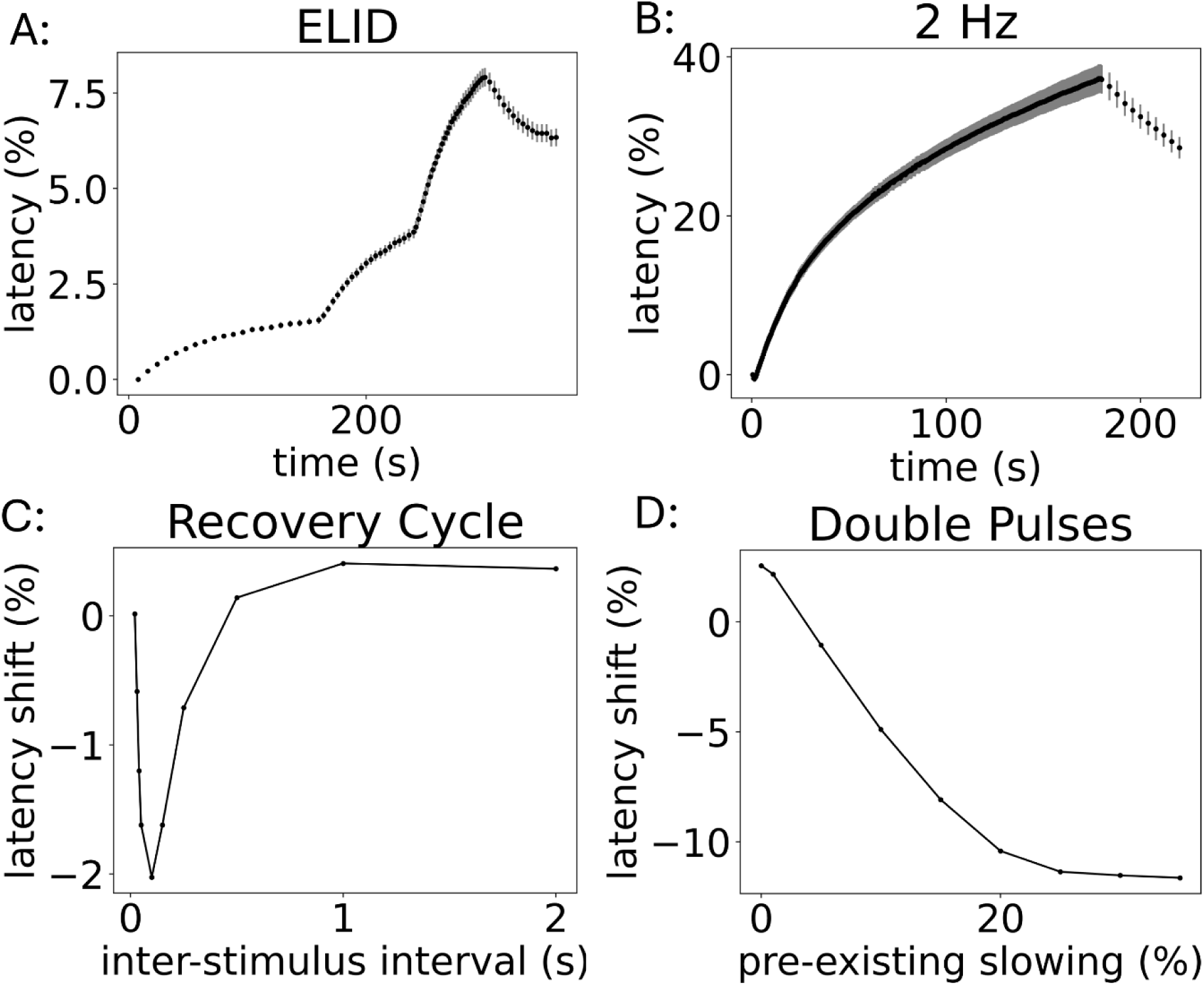
Overview of available data and stimulation protocols. A: Microneurography data from Namer’s lab for ELID protocol with mean and standard error of the mean. B: Microneurography data from Namer’s lab for 2 Hz protocol with mean and sem. C: Data from (Weidner et al., 2000) for recovery cycle protocol. The latency shift is plotted for each inter-stimulus interval. D: Data from (Weidner et al., 2002) for double pulse protocol.

The microneurography data obtained from these protocols were subsequently analyzed by extracting the latencies for all action potentials recorded during the ELID and 2 Hz stimulation. For each fiber, the latencies were normalized to the base latency obtained after the initial 2-minute pause. For further usage in the optimization process, the mean latency across all fibers was computed.

#### 2.3.3. Data from Literature

In addition to our original dataset, we incorporated findings from (Weidner et al., 2002, 2000), which focuses on small inter-stimulus intervals. The data was extracted directly from figures in these papers using PlotDigitizer (https://plotdigitizer.com/).

Two distinct protocols were employed in these studies:

1. Recovery Cycle: Regular stimulation pulses at ¼ Hz frequency were given until the latency stabilized. Subsequently, additional pulses were interposed at varying intervals defined as follows: 2s, 1s, 0.5s, 0.25s, 0.15s, 0.1s, 0.05s, 0.04s, 0.03s, 0.02s.
2. Double pulses: Two stimulation pulses were delivered with an interval of either 20 ms or 50 ms, with each pair repeated at frequencies ranging from 1/16 to 1 Hz.

The recovery cycle data were obtained from (Weidner et al., 2000), who examined 16 fibers. This data represents the mean latency shift of the base pulses in response to the extra stimulation pulses, plotted against the corresponding inter-stimulus interval (Figure 4C).

The double pulse data was obtained from (Weidner et al., 2002), Figure 4, who examined a total of 51 fibers. This data represents the mean latency shift, i.e. the difference in latency between the second and first pulse of the pair, plotted against the pre-existing activity state, defined as the relative latency of the preceding action potential (Figure 4D).

### 2.4. Parameter Optimization

The optimization of the model function was conducted for the linear and non-linear parts separately, each using stimulation protocols containing inter-spike intervals (ISIs) in the respective range.

The linear part of the memory function using ISIs ≤ 0.5 s was optimized using the ELID and 2 Hz stimulation protocols. Both protocols exclusively contain frequencies within this range.

The protocols were evaluated within the model, and the mean squared error of the simulated latencies and the corresponding experimental data was minimized.

Once the parameters for the linear component were established, the exponential part using ISIs > 0.5 s was optimized. This phase utilized data from the Recovery Cycle (RC) protocol, comparing simulated latency shifts against experimental data by calculating mean squared errors across all shifts to assess model performance effectively. Figure 5 displays an overview of the optimization and validation process.

**Figure 5:**
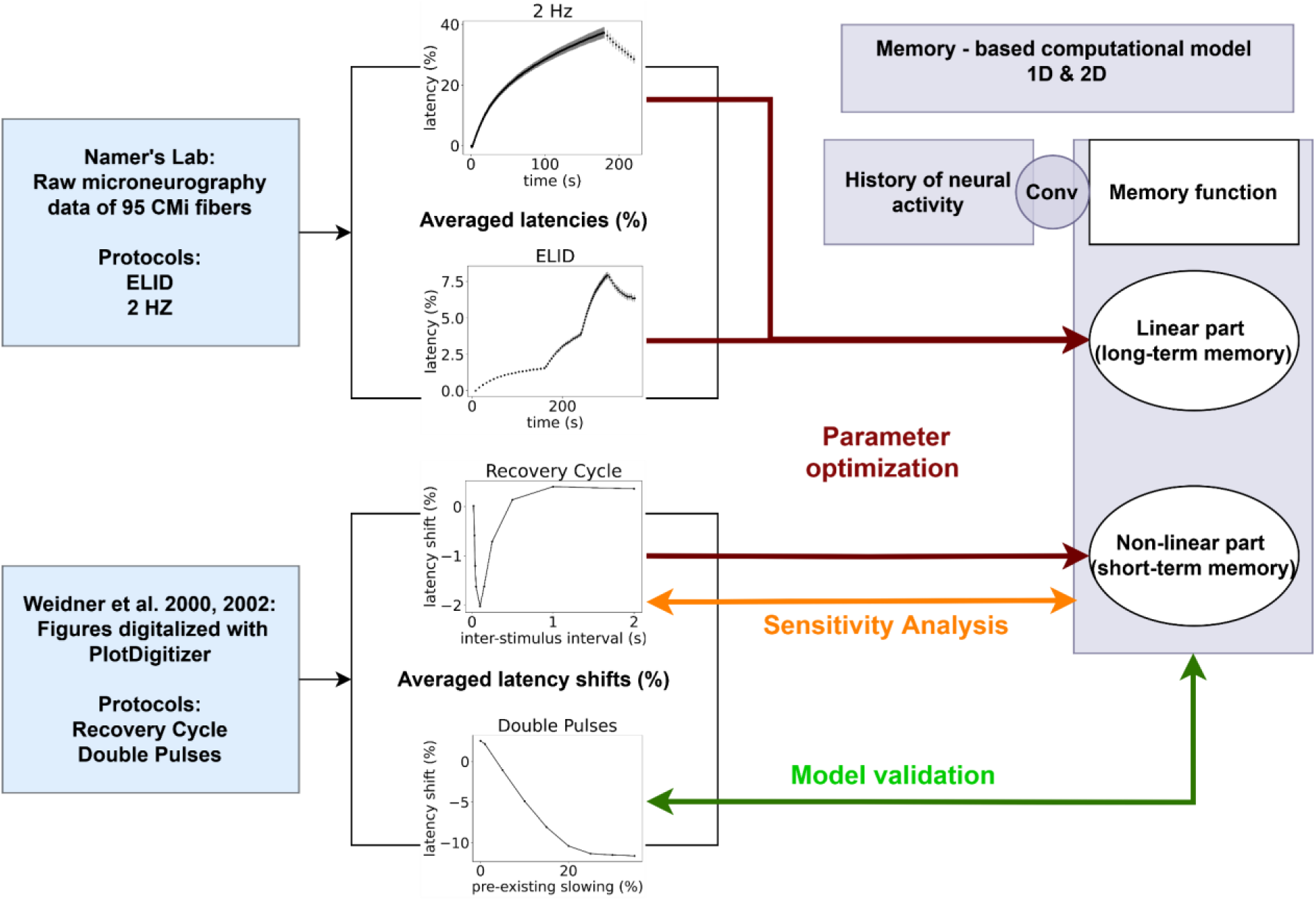
Overview of Optimization Process: The linear part of the memory function, representing long term axonal memory, was optimized using microneurography data from Namer’s lab that contained stimulation protocols with inter-stimulus intervals larger than 0.5 seconds. The non-linear memory function, representing short term memory, was optimized using data from Weidner, 2000, 2002, with stimulation protocols that contained inter-stimulus intervals below 0.5 seconds. This data was further used for model validation and sensitivity analysis.

#### 2.4.1. Optimization Algorithm

For parameter optimization, a Particle Swarm Optimization (PSO) algorithm (Sedighizadeh et al., 2021) was used, which is a computational technique inspired by the social behavior of organisms, like swarms of birds or schools of fish.

The optimization process begins by randomly initializing the positions of 100 particles, where each particle represents a set of parameters for the memory function. These positions are constrained within predefined bounds, ranging from 0 up to 50, depending on the parameter, which are chosen based on the expected range of parameter values. The selection of bounds ensures the optimization process focuses on realistic solutions while allowing sufficient exploration of the parameter space.

For each particle, the model’s performance is evaluated by calculating a cost function, which quantifies the differences between the model’s predicted latencies and the experimental latencies. The cost function is computed as the sum of the squared differences between these values. This approach provides a robust measure of how well the current parameter set aligns with the experimental data. The computation of all 100 particles was done in parallel on the high-performance cluster of the RWTH Aachen.

The positions of the particles are updated iteratively using the Generalized Particle Swarm Optimization (GEPSO) algorithm (Sedighizadeh et al., 2021). This algorithm introduces enhancements to the standard PSO by incorporating additional factors into the position updates. Specifically, each particle’s position is adjusted based on the best position from a random particle, a random velocity component, the particle’s current velocity, the particle’s personal best position and the global best position of all particles in the swarm.

Compared to standard PSO, GEPSO increases the diversity of particle movements and reduces the risk of premature convergence by getting stuck in a local minimum. These make it suitable to find solutions even in complex parameter spaces.

The optimization process continues for a fixed number of iterations, which in this study was set to 1000. This stopping criterion balances computational efficiency with the need for sufficient iterations to achieve convergence.

To improve the efficiency of the optimization, a two-stage approach was employed. First, the linear component of the memory function was optimized using larger frequencies of stimulation pulses. For this optimization, the recorded microneurography data for the ELID protocol (frequencies ranging from 1/8 Hz to ½ Hz) and the 2 Hz protocol were applied. Once the linear part was optimized, the nonlinear component was refined using data from the recovery cycle protocol, which uses frequencies up to 66 Hz (inter-stimulus interval of 15 ms). Nonlinear effects only occur during shorter intervals, necessitating a distinct protocol to accurately capture these dynamics. Further, this stepwise approach ensures that the simpler aspects of the model are fine-tuned before addressing the more complex, nonlinear dynamics.

### 2.5. Validation

The double pulse protocol and modified recovery cycle protocols with different background frequencies and inter-stimulus intervals were used to validate the model. For the double pulse protocol, the mean squared error of the latency shifts was calculated to assess the agreement between the model’s predictions and the experimental data. For the modified recovery cycle protocols, no experimental data was available for direct comparison.

To evaluate the generalizability and predictive performance of the computational model, we validated it using stimulation protocols that were not included during the optimization process. These protocols were selected to provide diverse input conditions and ensure the model’s ability to predict responses beyond those it was directly trained on. The stimulation protocols used encompass a wide range of pre-existing activity states and inter-stimulus intervals, to capture the diverse conditions typically encountered in microneurography experiments.

### 2.6. Additional Protocols for Sensitivity Analysis

In addition to the primary protocols, augmentary stimulation protocols were implemented as proof of concept, for which no existing data for comparison are available. These additional protocols were based on the recovery cycle protocol. The first set of protocols varied the inter-stimulus intervals of the extra pulses. One protocol introduced noise to the ISIs, resulting in slight variations from the original intervals. The other protocol utilized ISIs that fell between the original values, specifically: 1.5 s, 0.75 s, 0.375 s, 0.2 s, 0.125 s, 0.075 s, 0.045 s, 0.035 s, 0.025 s. The second set of augmentary protocols explored varying frequencies for the background pulses, ranging from 1/20 Hz to 1/2 Hz.

The implementation of augmentary stimulation protocols enhances the robustness and applicability of our computational model. By exploring variations in inter-stimulus intervals (ISIs) and background pulse frequencies, we aim to assess the model’s sensitivity to changes that may occur in real-world scenarios. Given that existing data for comparison are lacking, these additional protocols provide an opportunity to investigate the impact of different stimulation patterns on action potential dynamics.

### 2.7 Code Accessibility

The code used in this research is available to promote transparency and reproducibility of the results. The complete source code, along with instructions for installation and usage, can be accessed at the repository Digital-C-fiber (Maxion et al., 2025).

## Results

### 3.1 One-Dimensional Memory Function

#### 3.1.1 Parameter Optimization

The optimization of the one-dimensional memory function was conducted using Particle Swarm Optimization (PSO) in two steps. In the first step, the linear component of the memory function was optimized. The obtained parameter values were a=0.19 and b=−0.00095. In the second step, the parameters for the exponential component of the function were refined. The PSO algorithm yielded values of c=12, d=4.69, e=1.59 and f=0.33. The final memory function is illustrated in Figure 6.

**Figure 6:**
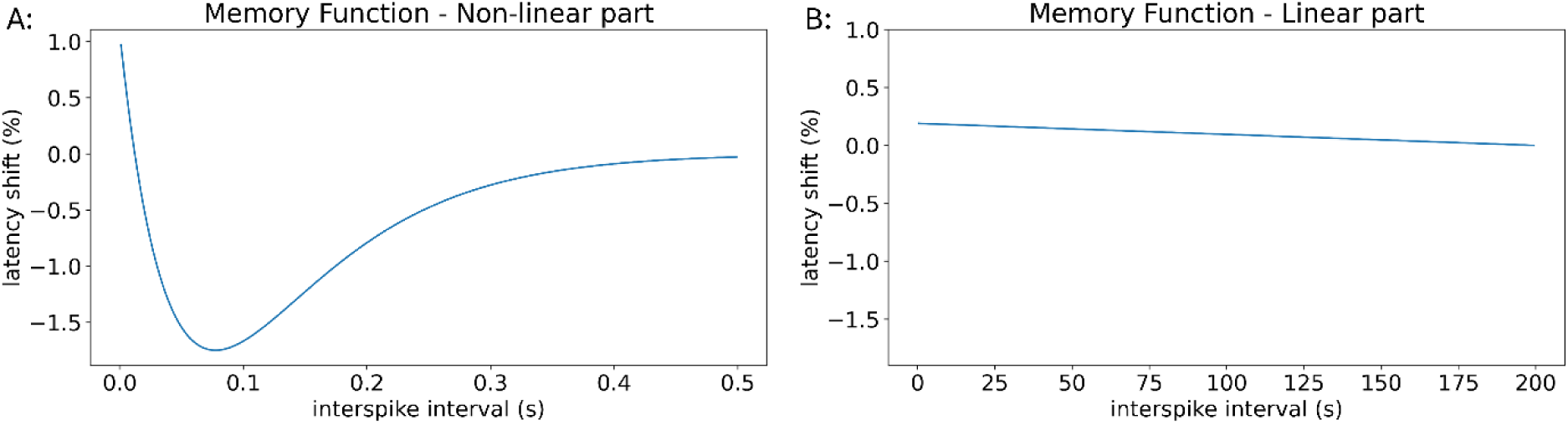
Illustration of the one-dimensional memory function with recovery cycle dynamics. A: The memory function for interspike intervals up to 0.5 s shows behaviour similar to recovery cycle dynamics with a supernormal phase. B: The memory function for interspike intervals larger than 0.5 s shows a linear behaviour.

#### 3.1.2 Model Performance

The performance of the convolution model utilizing the one-dimensional memory function was evaluated through its application to the ELID, 2 Hz and Recovery Cycle stimulation protocols. Figure 7 illustrates the simulated latencies compared to microneurography data from our own database (ELID and 2Hz) as well as data from (Weidner et al., 2002, 2000) (Recovery Cycle). The mean squared error was calculated for each case, yielding a value of 4.24 for 2 Hz stimulation and 0.41 for the ELID protocol (Table 1). For the recovery cycle protocol, the comparison between Weidner’s data and the simulated data resulted in a mean squared error of 0.09.

**Figure 7:**
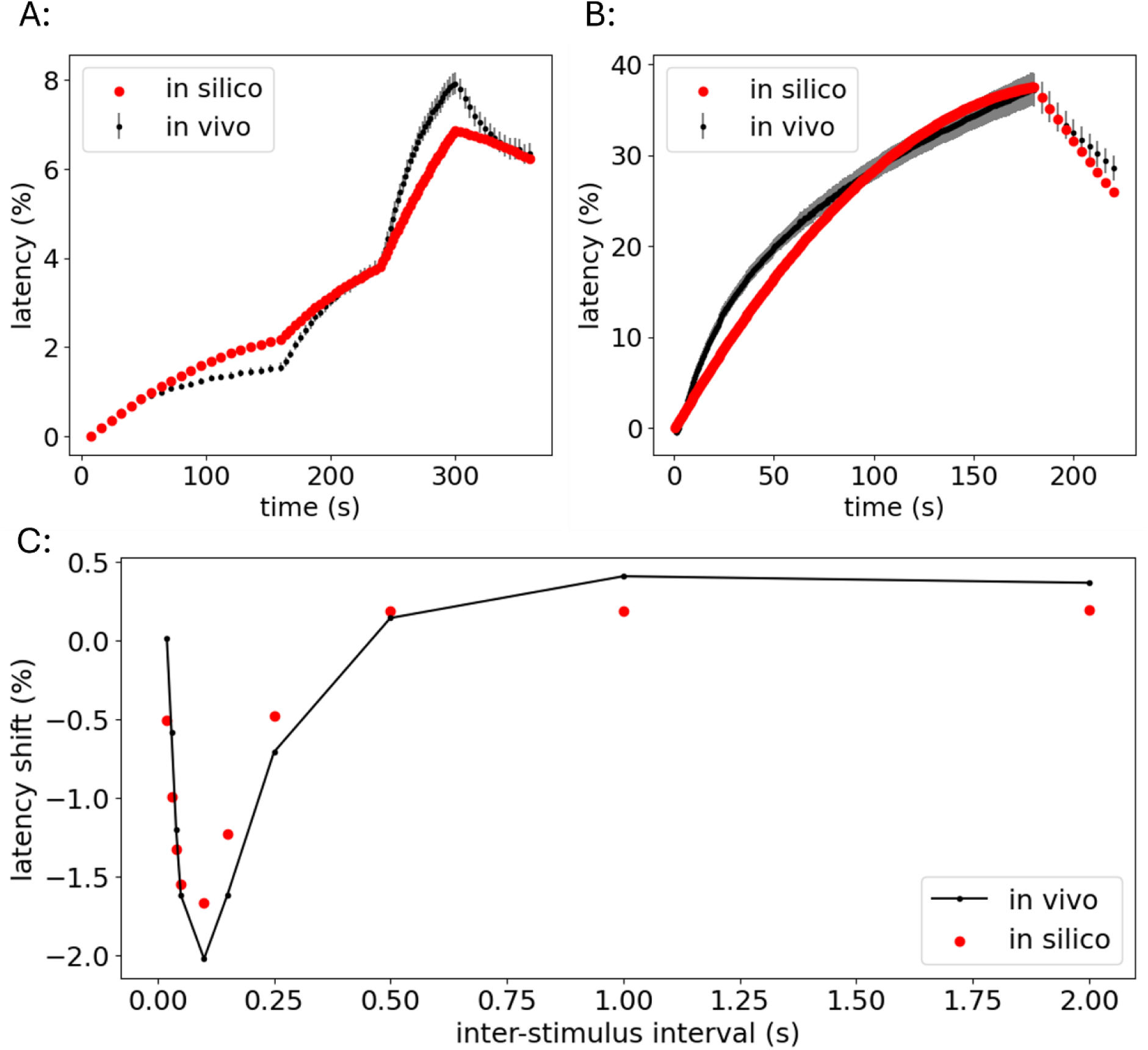
Model performance of the one-dimensional memory function. (A) and (B) Simulated latency increase expressed as a percentage (red) for the ELID and 2 Hz stimulation protocols, respectively. These are compared to latencies obtained from our own microneurography data (black), which displays the mean and standard error of the mean (SEM). (C) Relative latency shifts, also expressed as a percentage, calculated from the simulated latencies of the computational model (red) for the recovery cycle protocol. These shifts are compared to data from (Weidner et al., 2000) (black), showing the mean values.

**Table 1:** Mean Squared Errors (MSE) for the computational model using one-dimensional and two-dimensional memory functions across various stimulation protocols. Note that MSE values are only comparable between the different models for each protocol and should not be compared across different protocols.

|  | One-Dimensional | Two-Dimensional |
| --- | --- | --- |
| ELID | 0.41 | 0.41 |
| 2 Hz | 4.24 | 3.00 |
| Recovery Cycle | 0.09 | 0.027 |
| Double Pulses 20 ms | 12.20 | 1.87 |
| Double Pulses 50 ms | 51.14 | 17.45 |

#### 3.1.3 Validation

The model was validated using the double pulse protocol, which included inter-stimulus intervals of 20 ms and 50 ms at varying frequencies (Figure 8). As the frequency of the pulse pairs increases, resulting in greater pre-existing activity state, the latency difference between the two pulses remains constant. However, this behavior deviates substantially from what would occur in a real fiber, where experimental data indicates that increasing pre-existing activity state leads to a decrease in the latency shift between the pulses.

**Figure 8:**
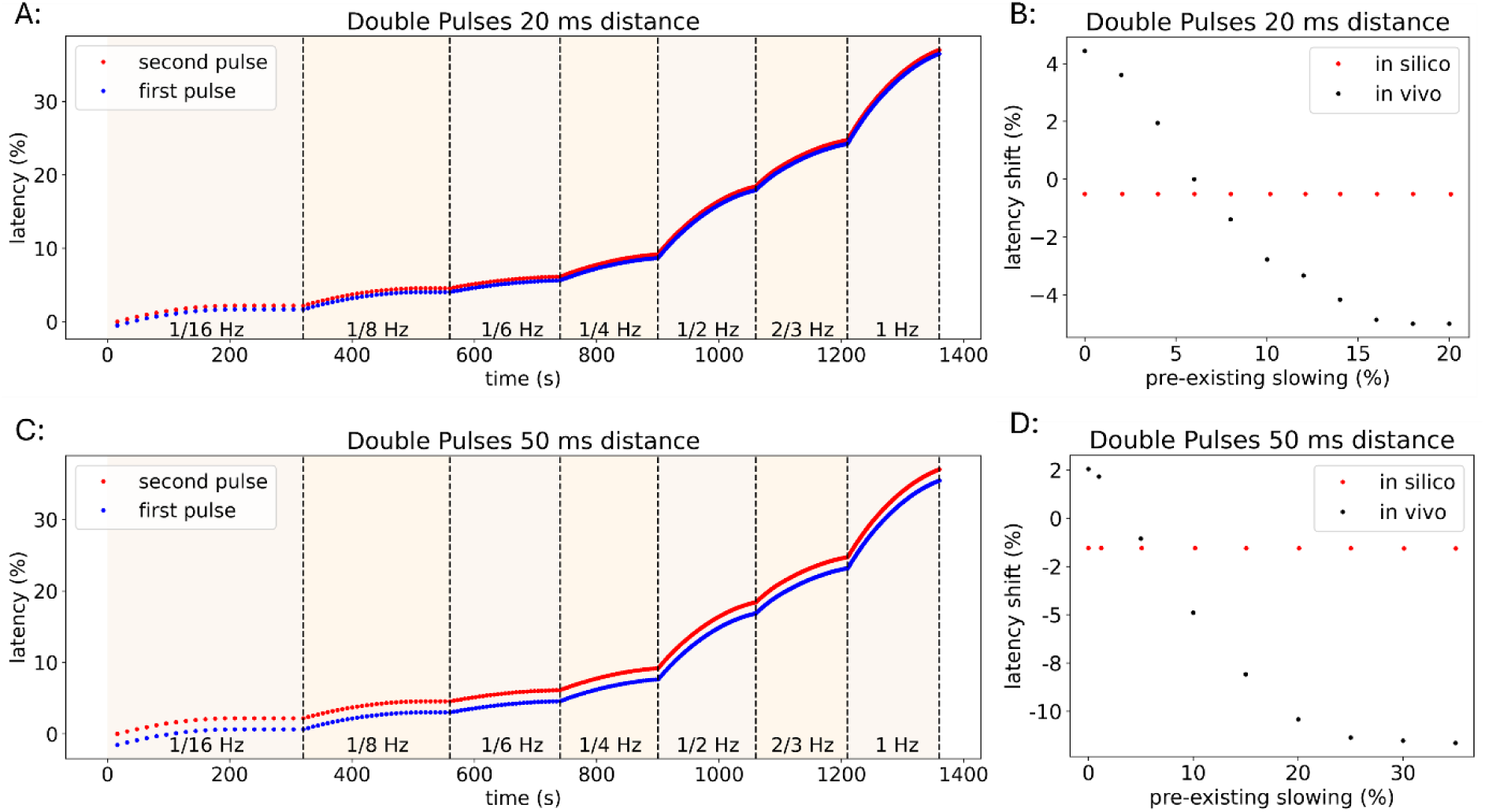
Results of the computational model for the double pulse protocols using the one-dimensional memory function. Two pulses are given at 20 ms or 50 ms distance at frequencies ranging from 1/16 Hz to 1 Hz. A + C: Relative latency over the time of the stimulation. B + D: The pre-existing activity state, represented as the relative latency of the preceding action potential, is compared to the latency shift, defined as the difference between the preceding and the current action potential. The real data is taken from Weidner, 2002.

#### 3.1.4. Sensitivity Analysis

Additionally, the recovery cycle protocol was modified by using different frequencies for the background pulses while maintaining consistent ISIs, see Figure 9. This led to varying levels of pre-existing activity state (Figure 9A) but produced the same relative latency shift across all frequencies (Figure 9B). This outcome contrasts with what we would expect in a real fiber, where the supernormal phase of the recovery cycle depends on the pre-existing activity state and varies with different frequencies.

**Figure 9:**
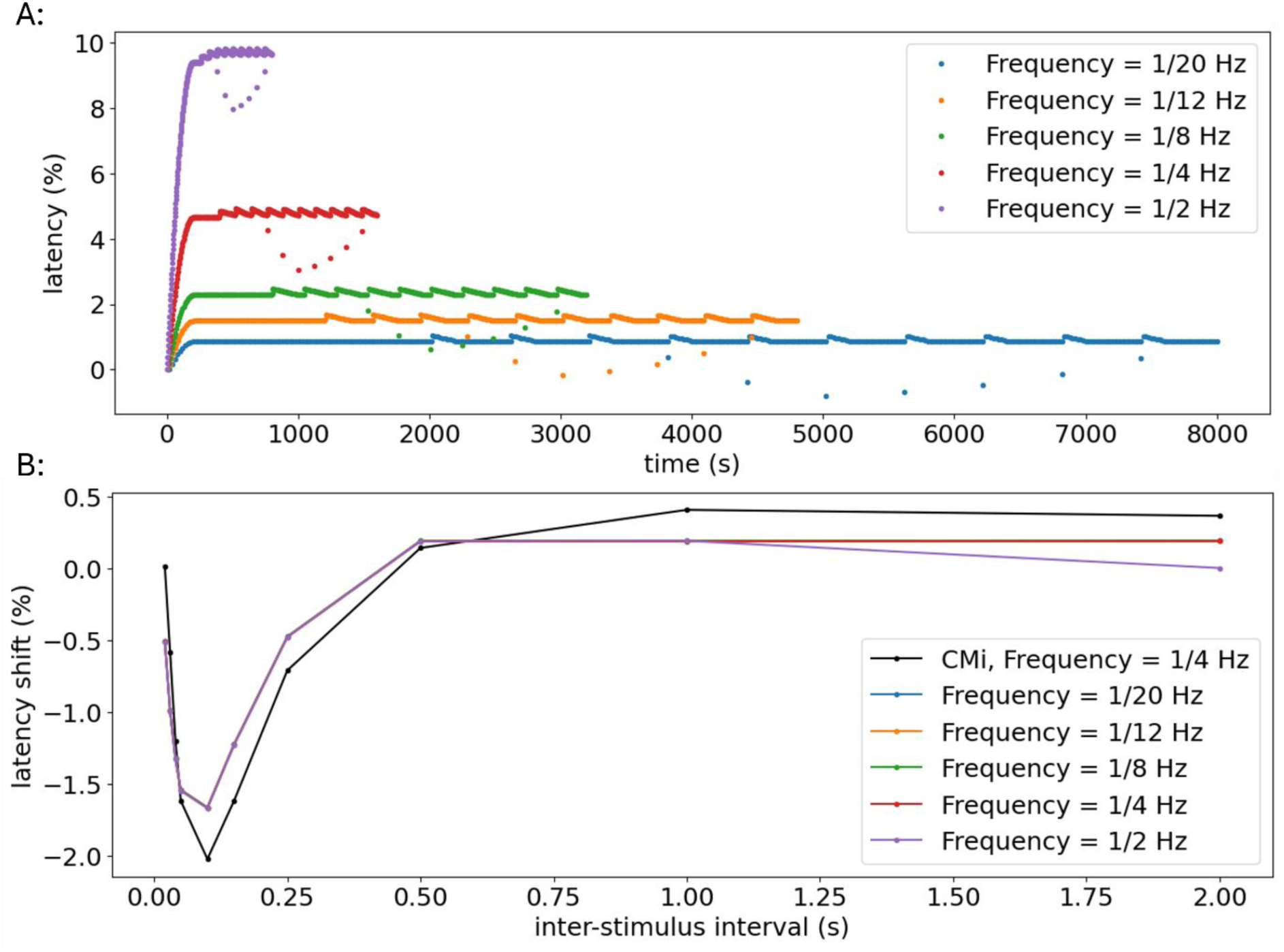
Results of the computational model for the recovery cycle protocol with varying frequencies of the background pulses using the one-dimensional memory function. A: Latency of the action potentials over the time of the stimulation. B: The inter-stimulus interval is shown compared to the relative latency shift induced by the extra pulses. In black the values of the real data from Weidner, 2000, are given for a background frequency of ¼ Hz.

This highlights the limitation of the one-dimensional model. Although the recovery cycle function captures the behaviour of the fiber under various ISIs, it does not account for the effects of pre-existing activity state. When the recovery cycle protocol is performed with a higher frequency of background pulses, the model is not able exhibit the expected more pronounced speeding for small ISIs. Instead, the latency shifts in the recovery cycle protocol remain the same regardless of the background frequency used. This discrepancy suggests that the current model lacks the ability to represent the full range of dynamic changes that occur with varying pre-existing activity state.

### 3.2 Two-Dimensional Memory Function

#### 3.2.1 Parameter Optimization

The parameters of the two-dimensional memory function (Equation 2) were optimized using Particle Swarm Optimization (PSO), following the same methodology applied to the one-dimensional function. This optimization was conducted in two steps: first, we focused on optimizing the linear component, resulting in values of a=0.22, b=0.0014, and c=0. In the second step, we refined parameters for the non-linear part of the function, yielding values of d = 21.800, e1 = 0.105, e2 = 0.0, f = 7.530, g = 0.553, h = 2.851, q = 3.485. The resulting memory function is illustrated in Figure 10.

**Figure 10:**
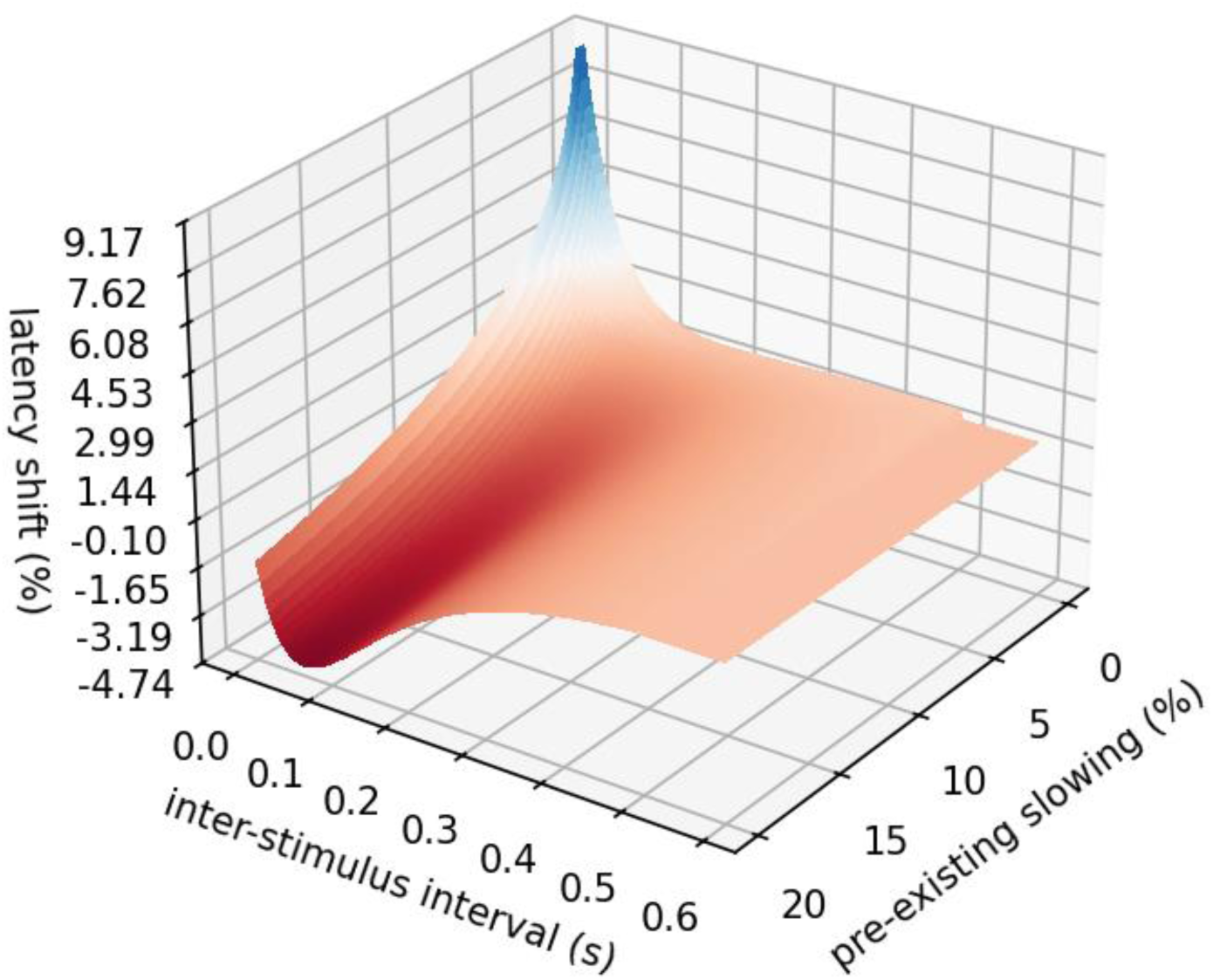
Illustration of the two-dimensional exponential memory function, predicting the latency shift of an action potential given the inter-stimulus interval and the pre-existing activity state.

In Figure 11 the optimization process is illustrated, showing the scores of all evaluated particles. The distribution of evaluated particles revealed clusters of high-performing parameter sets, suggesting specific regions of parameter space that contribute most effectively to model performance.

**Figure 11:**
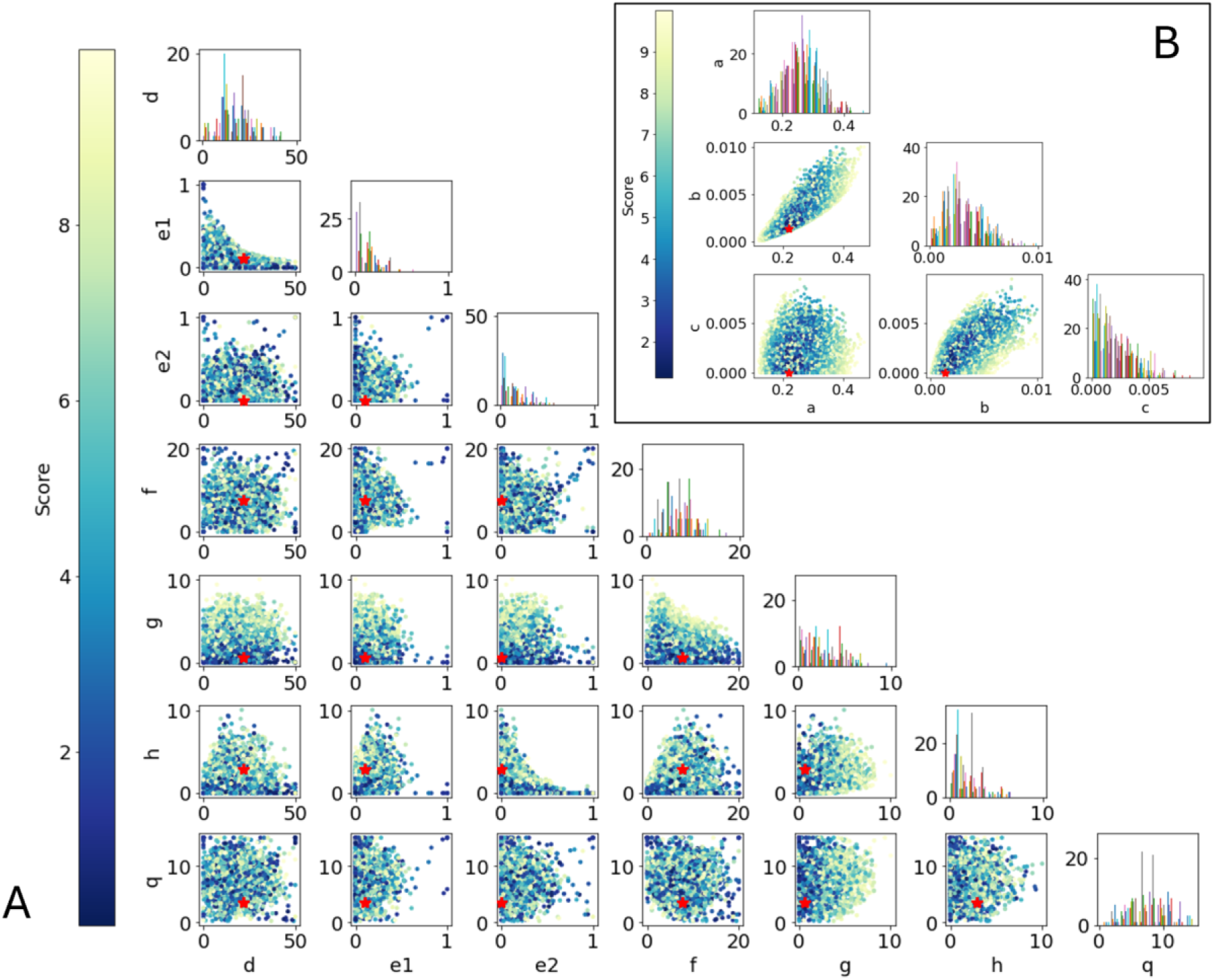
Results of the parameter optimization using a particle swarm algorithm. Each dot represents one particle, i.e. one evaluation of the model for a specific parameter set. The color indicates the fit/score of the model with lower values indicating a better fit. Only particles with scores below 10 are shown. The red star marks the best particle. The histograms show how many particles were evaluated in each parameter region. A: Parameter optimization for the non-linear component of the memory function. B: Parameter optimization for the linear component of the memory function.

#### 3.1.2 Model Performance

To assess the performance of the model, we evaluated its predictive accuracy across different protocols. The mean squared error for the ELID protocol is 0.41, while for the 2 Hz protocol, it stands at 3.00 (Table 1). The memory function captures the behavior of the post-excitatory effects well and preserves defining features, for example, an exponential behavior for low pre-existing activity state and the characteristic dip for low ISIs. The model is able to replicate the recovery cycle well with a mean squared error of 0.027 (Figure 12).

**Figure 12:**
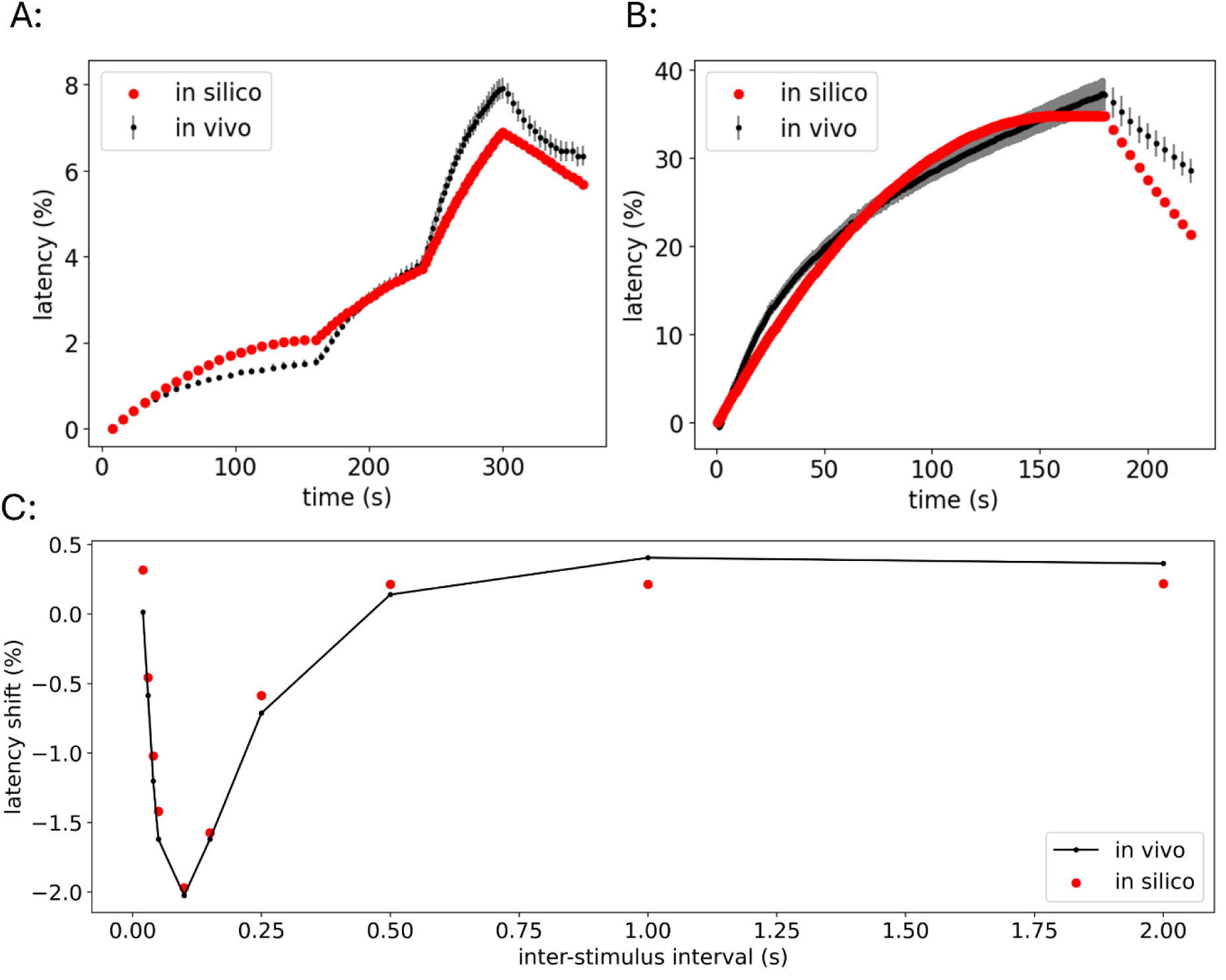
(A) and (B) Results for the ELID and 2 Hz protocols for the computational model using a two-dimensional linear memory function. (C) Results of the computational model (red) for the recovery cycle protocol compared to microneurography data (black) that shows the mean for the non-linear one-dimensional memory function.

#### 3.2.3 Validation

The last protocol used for validation is the double pulse protocol, where two pulses at 20 ms or 50 ms distance are given at different frequencies ranging from 1/16 Hz to 1 Hz (Figure 13). For higher frequencies, the second pulse shows activity-dependent speeding, while for lower frequencies, the second pulse is relatively slower than the first pulse. The computational model was able to capture the dynamics well.

**Figure 13:**
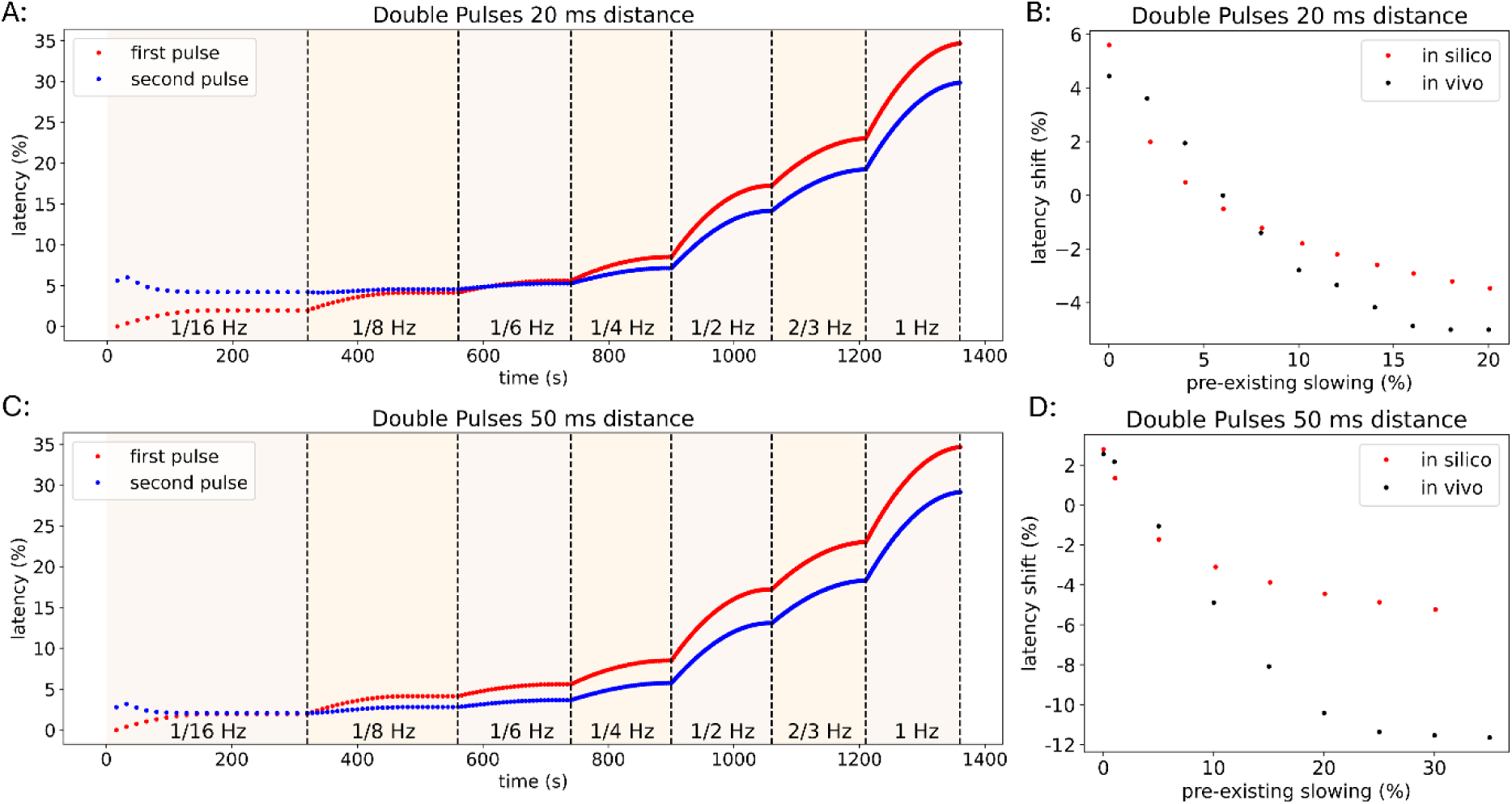
Results of the computational model for the double pulse protocols using the second two-dimensional memory function. Two pulses are given at 20 ms or 50 ms distance at frequencies ranging from 1/16 Hz to 1 Hz. A + B: Relative latency over the time of the stimulation. B + D: The pre-existing activity state, represented as the relative latency of the preceding action potential, is compared to the latency shift, defined as the difference between the preceding and the current action potential. The in vivo data is from (Weidner et al., 2002).

#### 3.2.4. Sensitivity Analysis

Further, the recovery cycle protocol was adapted with different background frequencies that induce a different level of pre-existing activity state (Figure 14). The background frequencies ranged from 1/20 Hz to ½ Hz. To account for variability in stabilization points across different frequencies, we initiated each simulation with 100 pulses at ¼ Hz before interposing additional pulses. A higher frequency of the background pulses induced a higher pre-existing activity state, which led to a more pronounced supernormal phase in the recovery cycle.

**Figure 14:**
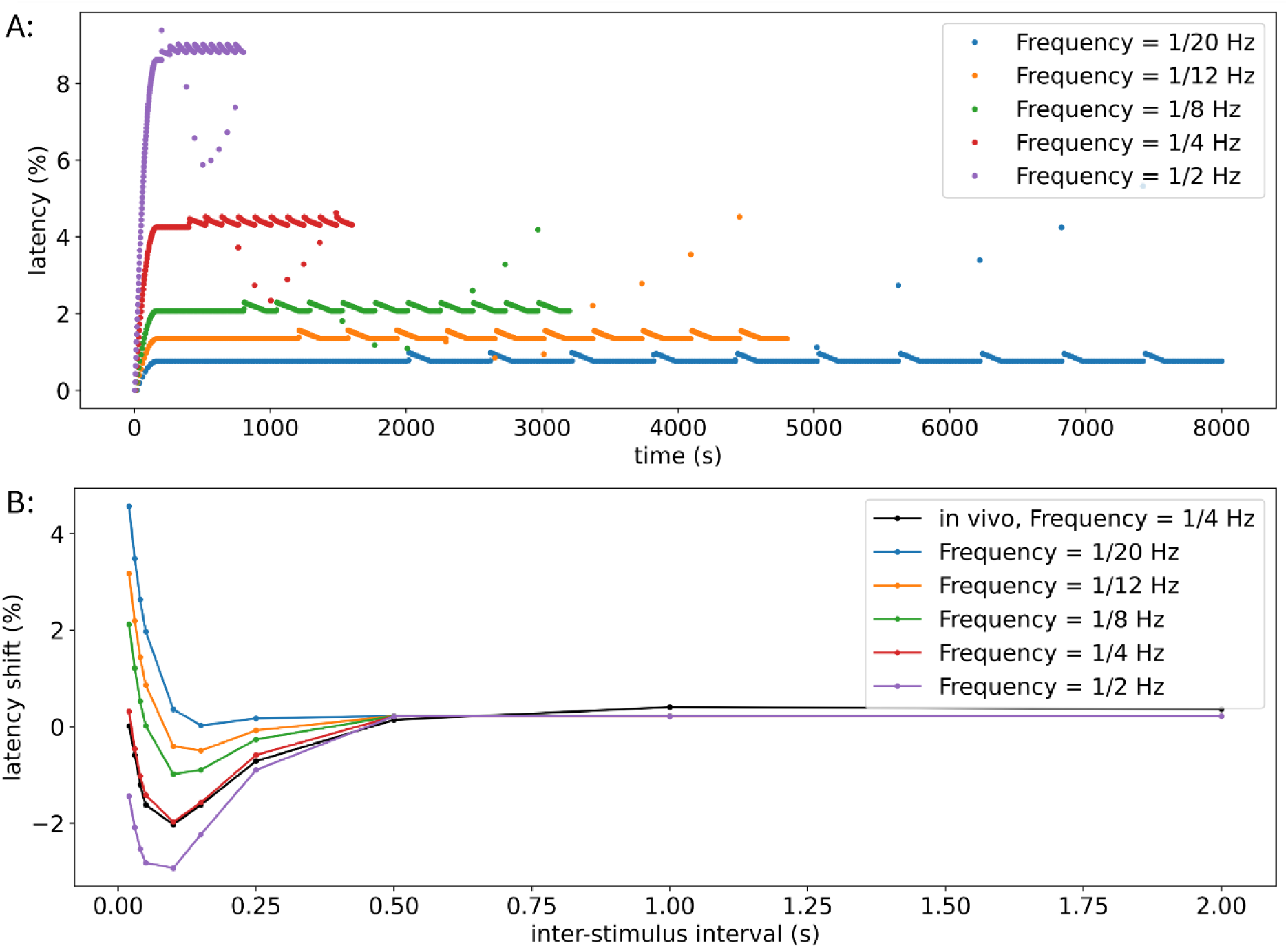
Results of the computational model for the recovery cycle protocol with varying frequencies of the background pulses using the two-dimensional memory function for sensitivity analysis. A: Latency of the action potentials over the time of the stimulation. B: The inter-stimulus interval is shown compared to the relative latency shift induced by the extra pulses. In black, the values of the real data from (Weidner et al., 2000), are given for a background frequency of ¼ Hz.

## 4. Discussion

We developed an innovative computational model for axonal action potential conduction based on the memory function of an axon. This model convolves a memory function with a signal function that contains the action potential initiation times. It efficiently predicts the conduction velocity of unmyelinated axons and thus opens the possibility for repetitive use in long axons, online applications, or parallel assessments of synchronicity.

In this project, we explored two potential memory functions: a one-dimensional function that is dependent on the inter-stimulus intervals (ISIs) and a two-dimensional function that is additionally dependent on the pre-existing activity state (PA). Both memory functions have in common that they include a linear component for larger ISIs and a non-linear component for shorter ISIs.

The model is validated using stimulation protocols that were not used for optimization and were chosen to cover a wide range of ISIs and a variety of pre-existing activity states to cover the whole parameter space.

### One-dimensional memory function

The one-dimensional memory function only depends on the ISI and is defined as a piecewise function with different dynamics for ISIs below and above 0.5 seconds. When action potentials are elicited with an ISI greater than 0.5 seconds, only activity-dependent slowing (ADS) of conduction velocity is observed. In contrast, for ISIs below 0.5 seconds, additional effects related to the recovery cycle can occur, leading to reduced ADS or even speeding of conduction velocity. This model shows a low mean squared error (MSE) for the ELID and 2 Hz protocols, see Table 1, which utilize stimulation frequencies at or below 2 Hz, and thus evoke mainly ADS correlating to the number of preceding action potentials. The main reason for ADS is thought to be intracellular sodium accumulation (Tigerholm et al., 2014), which constantly accumulates with each action potential, thereby generating most likely a linear effect. Thus, by modeling ADS with our approach we can also conclude on possible molecular processes underlying certain biological phenomena.

Stimulation protocols that include smaller ISIs, such as the recovery cycle protocol and the double pulse protocol, are reproduced utilizing additionally the non-linear component of the function, indicating that nonlinear mechanisms play an important role in these fast dynamics. This could be due to a combination of different processes, for example, sodium channel inactivation dependent on the membrane potential in combination with activity dependent hyperpolarization (Bostock et al., 2003; Bucher and Goaillard, 2011) and changes of intra-axonal and extra-axonal ionic concentrations (Tigerholm et al., 2015).

During the recovery cycle protocol, the model could replicate the supernormal phase, which plays a critical role in the dynamics of nerve fibers and can be altered in patients (Krishnan and Kiernan, 2005; Namer et al., 2015b). It can enhance the contrast of high and low activity in nerve fibers by modulating their timing, delaying them when prior activity is minimal and accelerating them when prior activity is substantial, resulting in a bursting pattern at the spinal cord (Bucher and Goaillard, 2011). Additionally, the model demonstrated robustness even when random variations were introduced to the existing recovery cycle protocol (Figure S1), indicating its reliability under various conditions.

However, the one-dimensional memory function does not account for pre-existing activity state, leading to the same results regardless of the amount of pre-existing activity, as seen in the results for the double pulse protocol (Figure 8) and the recovery cycle protocol with different background stimulation (Figure 9).

### Two-dimensional memory function

To solve the limitations of the one-dimensional memory function, the two-dimensional memory function includes the pre-existing activity state as an additional variable. Defined as a piecewise function, it exhibits different dynamics for ISIs below and above 0.5 seconds, with a two-dimensional linear function applied for larger ISIs. Parameter optimization revealed that the pre-existing activity state could be set to zero, indicating that for ISIs greater than 0.5 seconds, our model can accurately predict latencies using only the ISI of stimulation pulses, without explicitly considering prior activities, since ISIs inherently contain information about previous activity.

The component of the memory function for small ISIs consists of two exponential components subtracted from each other. Additionally, the second exponential term is multiplied by the logarithmic term and thus balances its impact. The resulting memory function resembles closely the function from (Weidner et al., 2002) and can reproduce a wide variety of stimulation protocols with different amounts of pre-existing activity and covering a wide range of ISIs accurately.

The MSE for the two-dimensional model was smaller than or equal to the MSE of the one-dimensional model for all protocols (Table 1). For increased background frequency in the recovery cycle protocol, which relates to more pre-existing activity, the supernormal phase of the recovery cycle increases, in accordance with the behaviour of real C-fibers (Bostock et al., 2003). Introducing noise in the recovery cycle protocol showed that the model maintained its accuracy, highlighting its robustness in varying conditions (Figure S2).

In conclusion, the model demonstrates robust agreement with experimental data and the optimized parameter set captures the essential physiological mechanisms underlying the observed behavior of unmyelinated axons. It captures key physiological phenomena, including different reactions to different stimulation frequencies and different levels of pre-existing activity state, both of which are critical for understanding the dynamic behavior of unmyelinated axons and thus the signal processing and memory.

### Comparison to existing computational models

Compared to existing models, this approach offers several distinct advantages. While Hodgkin-Huxley (HH) models of unmyelinated axons (He et al., 2020; Maxion et al., 2023; Tigerholm et al., 2014; Zhang et al., 2008) focus on incorporating biophysical details, our model prioritizes computational efficiency and interpretability. Despite its simplicity, it captures essential dynamics, including recovery cycles and pre-existing activity state. It can give insights into underlying molecular processes by correlating axonal dynamics with known biological processes, as discussed above.

One of the most significant advantages of our model is its fast computation time, making it suitable for real-time applications. The model’s computation times range from 2 to 100 ms, depending on the stimulation protocol, whereas a comparable HH model can take up to 20 minutes for the same protocols.

### Usability

The model effectively predicts spiking times of unmyelinated axons and its dynamics can be used to interpret underlying molecular mechanisms. Its supports simulations of longer axons or complex neural systems, implemented through an iterative loop, where each simulation’s output serves as the next input. This enables continuous modeling of neuronal activity along extended axonal pathways, in the case of sensory neurons, reaching up to the spinal cord, allowing for parallel simulations that assess synchronization or dyssynchronization crucial for sensory synaptic transmission and processing. This approach is relevant not only for pain but also for myelinated nerve fibers in other sensory systems, potentially aiding in prothesis development. For more complex neural systems involving multiple fibers with varying propagation properties, the memory function can be tailored to fit each individual fiber. Once these customized models are established, they can be interconnected by using the output from one neuron as the input for the subsequent neuron. This approach enables a comprehensive simulation of interactions within networks of interconnected neurons.

Furthermore, the model’s ability to represent both short and long ISIs highlights its versatility in addressing diverse experimental conditions, making it a valuable tool for both experimental analysis and hypothesis generation.

Additionally, it could assist during experiments by rapidly predicting spike times, both in real-time and during post-experiment analysis. In non-standard stimulation protocols, accurately predicting action potential latencies can be challenging, complicating the identification of spikes during the experiment. This computational model offers a rapid and efficient method to estimate where action potentials are likely to occur, aiding in their detection and analysis. In the future, these predictions might be incorporated into probabilistic decision-making layers for automatic spike detection and spike sorting for multifiber recordings.

Further it could be used to simulate stimulation protocols that cannot be practically or ethically performed e.g. during microneurography in human, for example, high-frequency protocols that cause pain in volunteers.

The model’s simplicity also makes it highly adaptable to other axonal types e.g. sympathetic fibers, which are involved in various pathologies, including vegetative nerve fibers controlling vital functions such as heart rate and blood pressure or axons of the central nervous system. By adjusting the memory function for patient data with abnormal nerve function, it may offer information about the mechanisms involved in pathological conditions.

These strengths position the model as a complementary tool to existing frameworks, particularly when rapid computation and flexibility are required.

### Limitations

Despite its strengths, the model has several limitations. First, defining an accurate memory function remains challenging, as it requires balancing simplicity and fidelity to experimental data. The optimization process, while effective, may be computationally intensive, particularly when integrating more complex memory functions. Additionally, the model does not explicitly incorporate information about specific ion channels, limiting its ability to offer mechanistic explanations for observed phenomena onto an indirect approach via dynamics.

## 5. Conclusion

This study presents a computational model that effectively captures the dynamics of axonal memory in unmyelinated axons, offering a systematic approach to understanding signal processing along unmyelinated axons. Among the two approaches tested, the two-dimensional memory function, depending on the inter-stimulus intervals and the pre-existing activity state, emerged as most accurate. It demonstrated robust performance across a variety of stimulation protocols, including protocols not used in the optimization process, while remaining computationally efficient.

The model’s utility lies in its ability to test predictions about axonal memory in a controlled and efficient manner, which is not feasible through experimental methods alone due to the significant time investment required and technical restrictions. By integrating predictions from the model with experimental observations, researchers can gain deeper insight into membrane processes shaping action potential propagation and their physiological and pathological changes.

One of the key advantages of the model is its computational efficiency, possibly enabling real-time applications, for example predicting action potential timing during axonal recordings such as microneurography experiments. This capability is particularly valuable for analyzing non-standard stimulation protocols or unstimulated activity where latency prediction remains challenging. Via this it could be a valuable addition to spike detection and sorting algorithms.

Moreover, its adaptability makes it a promising candidate for future enhancements, in particular adjustments to model different fiber types (e.g., unmyelinated and different myelinated axons) or to model diseased patient-specific abnormal axonal behaviour.

## Author contributions

AM: Formal analysis, Software, Visualization, Writing – original draft, Writing – review & editing

BN: Data acquisition, Data curation, Supervision, Writing – review & editing.

JT: Supervision, Writing – review & editing

EK: Conceptualization, Supervision, Writing – review & editing

## Acknowledgments

Computations were performed with computing resources granted by RWTH Aachen University under project rwth1541.

We thank all volunteers who participated. We thank Danxia Bao for excellent technical support.

## 7. Supplemental Material

**Figure S1:**
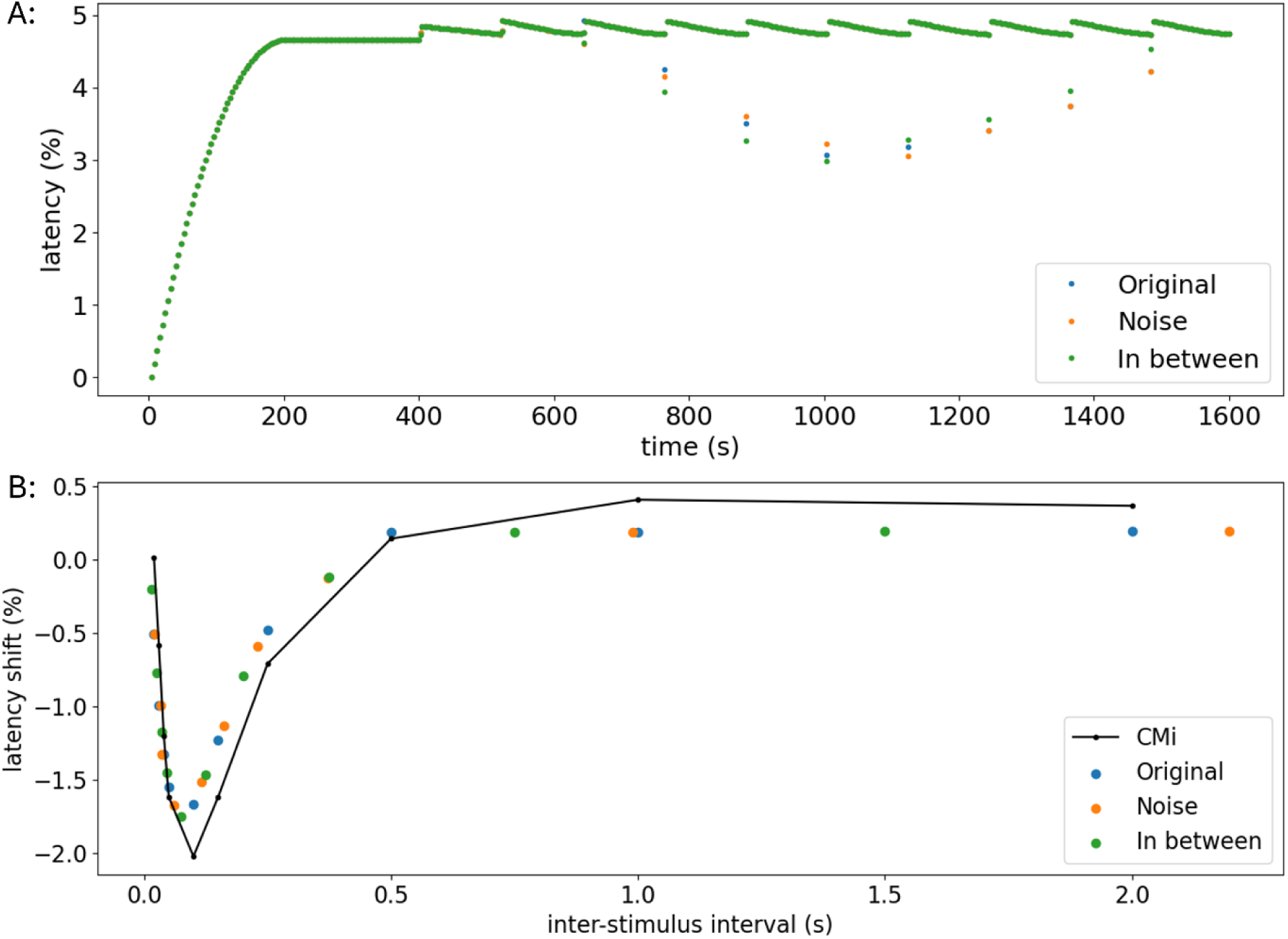
Results of the computational model for the recovery cycle protocol with varying distances between inter-stimulus pulses and a background frequency of ¼ Hz using the one-dimensional memory function. To demonstrate the robustness of the model, the recovery cycle protocol was evaluated under two additional conditions. To assess the model’s performance under conditions that mimic experimental variability, random noise was added to the additional pulses in the recovery cycle protocol. The model maintained a high degree of accuracy. The protocol was further tested using ISIs interpolated between the original timings. Again, the model exhibited a strong fit, indicating its ability to accurately predict recovery dynamics across a broader range of conditions. Blue: previously defined inter-stimulus intervals; Orange: Noise is added to the inter-stimulus intervals to get random variations; Green: The distance of the inter-stimulus intervals lies in between the previously defined ones. A: Latency of the action potentials over the time of the stimulation. B: The inter-stimulus interval is shown compared to the relative latency shift induced by the extra pulses. In black the values of the real data from Weidner, 2000, are shown.

**Figure S2:**
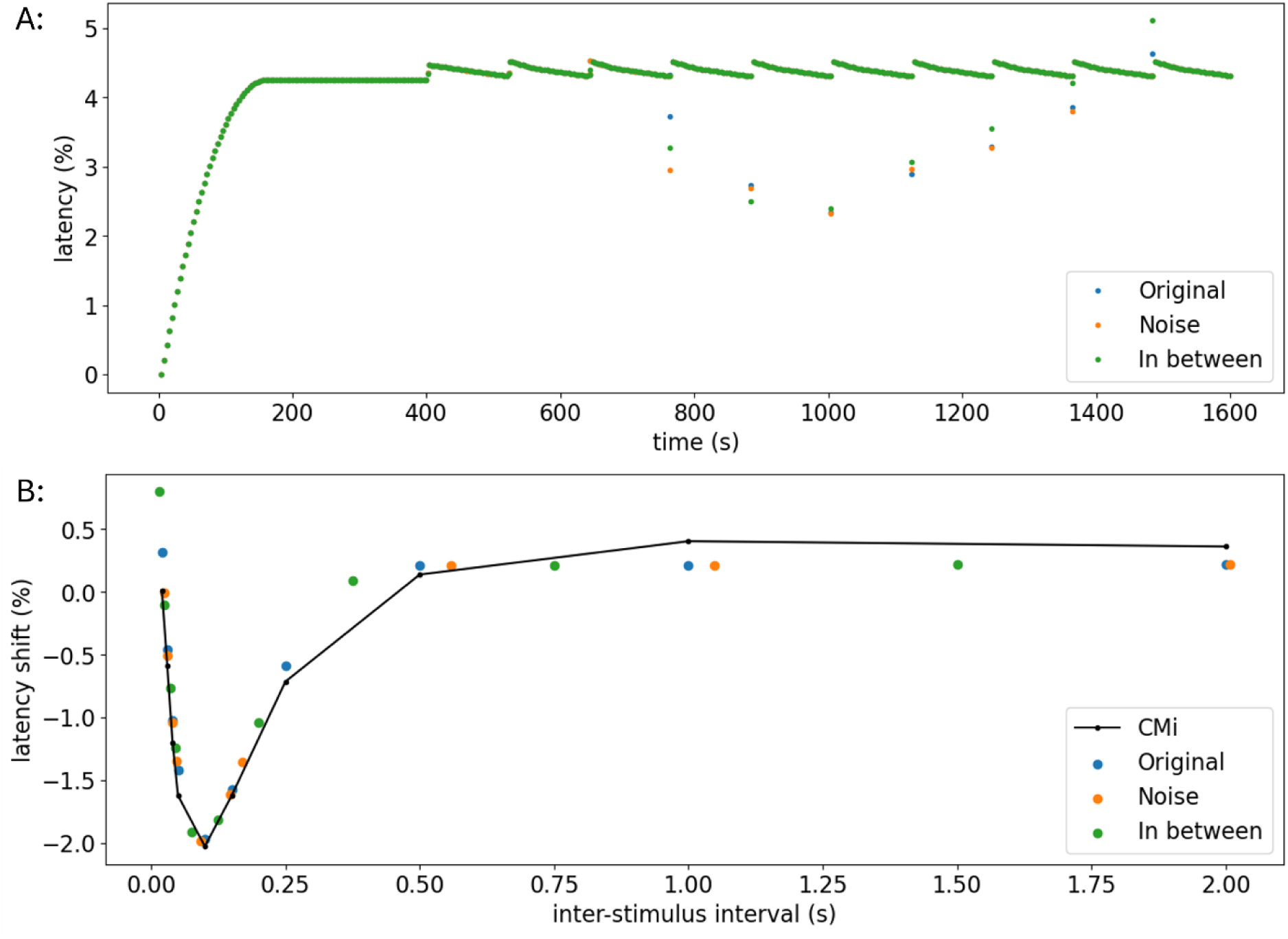
Results of the computational model for the recovery cycle protocol with varying distances between inter-stimulus pulses and a background frequency of ¼ Hz using the two-dimensional memory function. For the sensitivity analysis of the model using the two-dimensional memory function, we tested the recovery cycle protocol with modified ISIs. Both modified protocols could replicate the experimental data well. Blue: previously defined inter-stimulus intervals; Orange: Noise is added to the inter-stimulus intervals to get random variations; Green: The distance of the inter-stimulus intervals lies in between the previously defined ones. A: Latency of the action potentials over the time of the stimulation. B: The inter-stimulus interval is shown compared to the relative latency shift induced by the extra pulses. In black the values of the real data from Weidner, 2000, are shown.

## References

Ackerley, R., Watkins, R.H., 2018. Microneurography as a tool to study the function of individual C-fiber afferents in humans: responses from nociceptors, thermoreceptors, and mechanoreceptors. J. Neurophysiol. 120, 2834–2846. 10.1152/jn.00109.2018

Beeman, D., 2022. Hodgkin-Huxley Model, in: Encyclopedia of Computational Neuroscience. Springer, New York, NY, pp. 1627–1638. 10.1007/978-1-0716-1006-0_127

Bostock, H., Campero, M., Serra, J., Ochoa, J., 2003. Velocity recovery cycles of C fibres innervating human skin. J. Physiol. 553, 649–663. 10.1113/jphysiol.2003.046342

Bucher, D., Goaillard, J.-M., 2011. Beyond faithful conduction: short-term dynamics, neuromodulation, and long-term regulation of spike propagation in the axon. Prog. Neurobiol. 94, 307–346. 10.1016/j.pneurobio.2011.06.001

Carr, R., 2013. Nociceptors and Activity-Dependent Changes in Axonal Conduction Velocity, in: Encyclopedia of Pain. Springer, Berlin, Heidelberg, pp. 2244–2249. 10.1007/978-3-642-28753-4_4984

Dickie, A.C., McCormick, B., Lukito, V., Wilson, K.L., Torsney, C., 2017. Inflammatory Pain Reduces C Fiber Activity-Dependent Slowing in a Sex-Dependent Manner, Amplifying Nociceptive Input to the Spinal Cord. J. Neurosci. 37, 6488–6502. 10.1523/JNEUROSCI.3816-16.2017

Dubin, A.E., Patapoutian, A., 2010. Nociceptors: the sensors of the pain pathway. J. Clin. Invest. 120, 3760–3772. 10.1172/JCI42843

He, S., Yoshida, Y., Tripanpitak, K., Takamatsu, S., Huang, S.Y., Yu, W., 2020. A Simulation Study on Selective Stimulation of C-Fiber Nerves for Chronic Pain Relief. IEEE Access 8, 101648–101661. 10.1109/ACCESS.2020.2997964

Kist, A., Sagafos, D., Rush, A., Neacsu, C., Eberhardt, E., Schmidt, R., Lunden, L., Ørstavik, K., Kaluza, L., Meents, J., Zhang, Z., Carr, H., Salter, H., Malinowsky, D., Wollberg, P., Krupp, J., Kleggetveit, I., Schmelz, M., Jørum, E., Namer, B., 2016. SCN10A Mutation in a Patient with Erythromelalgia Enhances C-Fiber Activity Dependent Slowing. PloS One 11, e0161789. 10.1371/journal.pone.0161789

Kleggetveit, I.P., Namer, B., Schmidt, R., Helås, T., Rückel, M., Ørstavik, K., Schmelz, M., Jørum, E., 2012. High spontaneous activity of C-nociceptors in painful polyneuropathy. PAIN® 153, 2040–2047. 10.1016/j.pain.2012.05.017

Krishnan, A.V., Kiernan, M.C., 2005. Altered nerve excitability properties in established diabetic neuropathy. Brain 128, 1178–1187. 10.1093/brain/awh476

Kutafina, E., Troglio, A., de Col, R., Röhrig, R., Rossmanith, P., Namer, B., 2022. Decoding Neuropathic Pain: Can We Predict Fluctuations of Propagation Speed in Stimulated Peripheral Nerve? Front. Comput. Neurosci. 16. 10.3389/fncom.2022.899584

Maxion, A., Kutafina, E., Dohrn, M.F., Sacré, P., Lampert, A., Tigerholm, J., Namer, B., 2023. A modelling study to dissect the potential role of voltage-gated ion channels in activity-dependent conduction velocity changes as identified in small fiber neuropathy patients. Front. Comput. Neurosci. 17. 10.3389/fncom.2023.1265958

Maxion, A., Tigerholm, J., Namer, B., Kutafina, E., 2025. Modelling-of-Memory-in-Unmyelinated-Axons. 10.5281/zenodo.17182121

N. van Dalen, K., Slob, E., Schoemaker, C., 2013. Generalized minimum-phase relations for memory functions associated with wave phenomena. Geophys. J. Int. 195, 1620–1629. 10.1093/gji/ggt297

Namer, B., Lampert, A., 2025. Functional signatures of human somatosensory C fibers by microneurography. Pain. 10.1097/j.pain.0000000000003605

Namer, B., Ørstavik, K., Schmidt, R., Kleggetveit, I.-P., Weidner, C., Mørk, C., Kvernebo, M.S., Kvernebo, K., Salter, H., Carr, T.H., Segerdahl, M., Quiding, H., Waxman, S.G., Handwerker, H.O., Torebjörk, H.E., Jørum, E., Schmelz, M., 2015a. Specific changes in conduction velocity recovery cycles of single nociceptors in a patient with erythromelalgia with the I848T gain-of-function mutation of Nav1.7. PAIN 156, 1637. 10.1097/j.pain.0000000000000229

Namer, B., Ørstavik, K., Schmidt, R., Kleggetveit, I.-P., Weidner, C., Mørk, C., Kvernebo, M.S., Kvernebo, K., Salter, H., Carr, T.H., Segerdahl, M., Quiding, H., Waxman, S.G., Handwerker, H.O., Torebjörk, H.E., Jørum, E., Schmelz, M., 2015b. Specific changes in conduction velocity recovery cycles of single nociceptors in a patient with erythromelalgia with the I848T gain-of-function mutation of Nav1.7. Pain 156, 1637– 1646. 10.1097/j.pain.0000000000000229

Obreja, O., Ringkamp, M., Namer, B., Forsch, E., Klusch, A., Rukwied, R., Petersen, M., Schmelz, M., 2010. Patterns of activity-dependent conduction velocity changes differentiate classes of unmyelinated mechano-insensitive afferents including cold nociceptors, in pig and in human. PAIN 148, 59–69. 10.1016/j.pain.2009.10.006

Ørstavik, K., Namer, B., Schmidt, R., Schmelz, M., Hilliges, M., Weidner, C., Carr, R.W., Handwerker, H., Jørum, E., Torebjörk, H.E., 2006. Abnormal Function of C-Fibers in Patients with Diabetic Neuropathy. J. Neurosci. 26, 11287–11294. 10.1523/JNEUROSCI.2659-06.2006

Ørstavik, K., Weidner, C., Schmidt, R., Schmelz, M., Hilliges, M., Jørum, E., Handwerker, H., Torebjörk, E., 2003. Pathological C-fibres in patients with a chronic painful condition. Brain 126, 567–578. 10.1093/brain/awg060

Petersson, M.E., Obreja, O., Lampert, A., Carr, R.W., Schmelz, M., Fransén, E., 2014. Differential Axonal Conduction Patterns of Mechano-Sensitive and Mechano-Insensitive Nociceptors – A Combined Experimental and Modelling Study. PLoS ONE 9, e103556. 10.1371/journal.pone.0103556

Schmelz, M., Forster, C., Schmidt, R., Ringkamp, M., Handwerker, H.O., Torebjörk, H.E., 1995. Delayed responses to electrical stimuli reflect C-fiber responsiveness in human microneurography. Exp. Brain Res. 104, 331–336. 10.1007/BF00242018

Schmelz, M., Schmidt, R., 2010. Microneurographic single-unit recordings to assess receptive properties of afferent human C-fibers. Neurosci. Lett., Microneurography: a relevant tool for the understanding of Neuropathic Pain 470, 158–161. 10.1016/j.neulet.2009.05.064

Schmidt, R., Schmelz, M., Forster, C., Ringkamp, M., Torebjörk, E., Handwerker, H., 1995. Novel classes of responsive and unresponsive C nociceptors in human skin. J. Neurosci. Off. J. Soc. Neurosci. 15, 333–341. 10.1523/JNEUROSCI.15-01-00333.1995

Sedighizadeh, D., Masehian, E., Sedighizadeh, M., Akbaripour, H., 2021. GEPSO: A new generalized particle swarm optimization algorithm. Math. Comput. Simul. 179, 194–212. 10.1016/j.matcom.2020.08.013

Serra, J., Campero, M., Ochoa, J., Bostock, H., 1999. Activity-dependent slowing of conduction differentiates functional subtypes of C fibres innervating human skin. J. Physiol. 515 ( Pt 3), 799–811. 10.1111/j.1469-7793.1999.799ab.x

Soleng, A.F., Chiu, K., Raastad, M., 2003. Unmyelinated axons in the rat hippocampus hyperpolarize and activate an H current when spike frequency exceeds 1 Hz. J. Physiol. 552, 459–470. 10.1113/jphysiol.2003.048058

Tigerholm, J., Petersson, M.E., Obreja, O., Eberhardt, E., Namer, B., Weidner, C., Lampert, A., Carr, R.W., Schmelz, M., Fransén, E., 2015. C-fiber recovery cycle supernormality depends on ion concentration and ion channel permeability. Biophys. J. 108, 1057– 1071. 10.1016/j.bpj.2014.12.034

Tigerholm, J., Petersson, M.E., Obreja, O., Lampert, A., Carr, R., Schmelz, M., Fransén, E., 2014. Modeling activity-dependent changes of axonal spike conduction in primary afferent C-nociceptors. J. Neurophysiol. 111, 1721–1735. 10.1152/jn.00777.2012

Torebjörk, H.E., Hallin, R.G., 1974. Identification of afferent C units in intact human skin nerves. Brain Res. 67, 387–403. 10.1016/0006-8993(74)90489-2

Vallbo, Å.B., 2018. Microneurography: how it started and how it works. J. Neurophysiol. 120, 1415–1427. 10.1152/jn.00933.2017

Weidner, C., Schmelz, M., Schmidt, R., Hammarberg, B., Ørstavik, K., Hilliges, M., Torebjörk, H.E., Handwerker, H.O., 2002. Neural Signal Processing: The Underestimated Contribution of Peripheral Human C-Fibers. J. Neurosci. 22, 6704– 6712. 10.1523/JNEUROSCI.22-15-06704.2002

Weidner, C., Schmelz, M., Schmidt, R., Hansson, B., Handwerker, H.O., Torebjörk, H.E., 1999. Functional Attributes Discriminating Mechano-Insensitive and Mechano-Responsive C Nociceptors in Human Skin. J. Neurosci. 19, 10184–10190. 10.1523/JNEUROSCI.19-22-10184.1999

Weidner, C., Schmidt, R., Schmelz, M., Hilliges, M., Handwerker, H.O., Torebjörk, H.E., 2000. Time course of post-excitatory effects separates afferent human C fibre classes. J. Physiol. 527 Pt 1, 185–191. 10.1111/j.1469-7793.2000.00185.x

Westenbroek, R.E., Merrick, D.K., Catterall, W.A., 1989. Differential subcellular localization of the RI and RII Na+ channel subtypes in central neurons. Neuron 3, 695–704. 10.1016/0896-6273(89)90238-9

Zhang, J., Wang, J., Liu, Y., Hu, S., 2008. Simulation of Primary Afferent Synapses in Unmyelinated Nerve Fiber, in: 2008 International Conference on BioMedical Engineering and Informatics. Presented at the 2008 International Conference on BioMedical Engineering and Informatics, pp. 723–727. 10.1109/BMEI.2008.364

